# Neural mechanisms of attention, not expectation, govern spatial selection by probabilistic cueing

**DOI:** 10.1101/2024.12.29.629912

**Authors:** Sricharan Sunder, Kavya Rajendran, Mainak Biswas, Devarajan Sridharan

## Abstract

Spatial probabilistic “Posner” cueing is widely employed in studies of endogenous spatial attention. Such cueing guides attention by providing prior knowledge about the likely spatial location of task-relevant events. Yet, it has been compellingly argued that such spatial priors also elicit expectation effects, rendering Posner cueing unsuitable for measuring attentional effects in isolation. We address this debate by combining signal detection theory models of behavior with concurrent electrophysiological recordings, and directly compare Posner cueing effects with those of attention and expectation cueing. Participants performed two tasks: a dual cueing task, with orthogonal relevance (attention) and probability (expectation) cues, as well as a Posner cueing task. Relevance and probability cueing independently modulated distinct behavioral parameters – perceptual sensitivity and decisional criterion, respectively – whereas Posner cueing modulated both sensitivity and criterion. However, only sensitivity modulations by Posner cueing were correlated with those of relevance cueing. Criterion modulations by Posner cueing were uncorrelated with those of probability cueing. Both Posner and relevance cueing, but not probability cueing, modulated various neural markers of spatial attention, including steady-state visually evoked potential (SSVEP) amplitude and alpha-band (8-12 Hz) oscillation power. Representational similarity analysis and cue-label prediction with deep convolutional neural networks revealed dissociable underpinnings of relevance and probability cueing, and identified Posner cueing’s neural representations with those of relevance cueing. Our results address a long-standing debate in the attention literature and clearly demonstrate that spatial selection by probabilistic cueing is governed by neural mechanisms of attention, not expectation.

**Significance Statement:** How does our brain select relevant information for accomplishing task goals? “Posner cueing” is a popular choice for studying brain mechanisms of goal-driven attention in the lab. Posner cues guide attention by providing advance information about the likely location of task-relevant events. Yet, Posner cueing has been criticized because it engages brain mechanisms not only of attention but of event expectation, as well. We address this critique by employing state-of-the-art neural decoding approaches, including deep convolutional neural networks, and show that behavioral and neural underpinnings of Posner cueing closely match those of attention, not expectation. The results address a long-standing debate in the attention literature and validate Posner cueing as a reliable method for measuring attention’s effects in the brain.

## Introduction

Our senses face a barrage of information from the world around us. Selective spatial attention enables us to select and process information from task relevant locations while ignoring or suppressing information from irrelevant locations. While spatial attention can be engaged in various ways – such as by manipulating rewards (Luo & Maunsell, 2015, 2018; Sali et al., 2014) or task relevance (Hawkins et al., 1990; Wyart et al., 2012) – anticipating a relevant event at a particular location can also engage spatial attention. Spatial probabilistic cueing, widely employed in laboratory tasks, exploits precisely such event anticipation to engage selective spatial attention. Such a cue, also called a “Posner cue” (following Posner, 1980), is informative regarding the location at which an upcoming task-relevant event (e.g. change) is most likely to occur. The expectation of the event encourages participants to direct their attention to its impending location.

Despite its ubiquity, the use of spatial probabilistic “Posner” cueing for studying attention is controversial. For many years (Duncan, 1980; Posner, 1980), it has been argued that spatial probabilistic cueing recruits both spatial attention and spatial expectation mechanisms, the latter two being dissociable cognitive processes (Doricchi et al., 2010; Kok, Rahnev, et al., 2012; Wyart et al., 2012). Despite this caveat, spatial probabilistic cueing has been the paradigm of choice for numerous studies characterizing behavioral and neural underpinnings of spatial attention, in humans (Geng & Behrmann, 2005; Giordano et al., 2009; Mangun & Hillyard, 1990; Sperling & Melchner, 1978), non-human primates (Ciaramitaro et al., 2001; M. R. Cohen & Maunsell, 2009; Cook & Maunsell, 2002; Raffi & Siegel, 2005; Ruff & Cohen, 2017), and even in non-primate animals (Giordano et al., 2009; Quest et al., 2022; Speed & Haider, 2021; Sridharan et al., 2013). Here, we seek to tease apart component cognitive processes, and neural mechanisms, engaged by spatial probabilistic Posner cueing. Specifically, we asked whether Posner cueing engages endogenous spatial attention, spatial expectation, or a combination of both processes. To answer this question, we examined evidence from literature regarding distinct behavioral and neural consequences of attention and expectation, respectively.

Behavioral studies seeking to find distinct effects of attention and expectation have reported mixed results. Studies investigating the effects of attention and expectation on reaction times (RTs) reported that endogenous attention produced faster RTs (Pham et al., 2018; Yantis & Jonides, 1984), but found mixed effects of expectation on RTs (Doricchi et al., 2010; Zuanazzi & Noppeney, 2019). A few studies have examined the effects of attention and expectation on the fidelity of sensory information processing (sensitivity) versus decision-making (criterion) with signal detection theory (SDT) (Green & Swets, 1974; Macmillan & Creelman, 2005). For example, Wyart et al (Wyart et al., 2012) showed that attention enhances perceptual sensitivity without producing systematic effects on criterion. Yet, other studies have suggested that attention produces dissociable effects on sensitivity and criterion (Luo & Maunsell, 2015, 2018). On the other hand, converging evidence from multiple studies indicates that manipulating expectation, by altering signal priors, produces a selective modulation of criterion but not sensitivity (Bang & Rahnev, 2017; Tarasi et al., 2022; Wyart et al., 2012; Zhou et al., 2021). By contrast, other studies have suggested that expectation for features produces perceptual benefits (Kok et al., 2017; Kok, Jehee, et al., 2012; Thomas et al., 2022). Here we asked whether Posner cueing would produce dissociable behavioral effects on sensitivity and criterion, mediated by distinct attention and expectation mechanisms, respectively.

Similarly, whether neural mechanisms that mediate attention and expectation are entirely dissociable or overlap partially, is an actively researched question (Doricchi et al., 2010; Gordon et al., 2019; Rungratsameetaweemana et al., 2018). Electrophysiological correlates of these cognitive processes have been explored extensively with scalp electroencephalography (EEG) in humans, with previous studies investigating both stimulus-driven steady-state visually evoked potentials (SSVEPs) (Mangun & Hillyard, 1990) as well as intrinsic alpha-band (8-12Hz) oscillations (Clayton et al., 2018). For example, SSVEP amplitude, a readout of early visual processing, increases significantly at the attended location (Morgan et al., 1996; Müller et al., 1998). Conversely, a few studies have suggested that expectation does not modulate visual information processing (Bang & Rahnev, 2017), although the effect of spatial expectation on SSVEP amplitudes remains to be systematically explored. Conversely, attention is widely reported to suppress alpha-band power in parieto-occipital channels contralateral to the attended stimulus location (Foxe & Snyder, 2011; Rihs et al., 2007; Thut et al., 2006) (but see Zhou et al, 2021). On the other hand, recent studies show that spatial expectation modulates both pre-stimulus alpha power (Tarasi et al., 2022) and alpha phase (Sherman et al., 2016). Based on this literature, we asked whether neural correlates of Posner cueing would overlap with established neural signatures of attention, expectation, or both.

Recent studies have sought to link neural correlates of attention – SSVEP and alpha power modulations – to behavioral modulations of sensitivity and criterion, respectively. For instance, a recent study employing a closed-loop paradigm (Chinchani et al., 2022) reported that visual presentation locked to relatively high SSVEP power states contralateral to the attended location yielded higher perceptual sensitivities for the target. This study concluded that instantaneous SSVEP power fluctuations may reflect a neural marker of visual sensitivity modulation by endogenous attention. By contrast, no study, to our knowledge, has systematically explored the link between SSVEP power and criterion modulations. Similarly, a few studies (Iemi et al., 2017; Limbach & Corballis, 2016; Samaha et al., 2020; Tarasi et al., 2022) reported that pre-stimulus alpha power modulation predicted bias (criterion) in the participant’s subsequent response. By contrast another recent study (Zhou et al., 2021) observed that although expectation modulated behavioral criteria, these effects were not correlated with pre-stimulus alpha power. Given these links, we asked whether Posner cueing effects on sensitivity and criterion would map on to distinct electrophysiological signatures – SSVEP and alpha-band power modulations, respectively.

To address each of these questions, we tested the same group of participants on two parallel task designs: a dual cueing task and a Posner cueing task (Fig. 1). The dual cueing task involved change detection with orthogonal attention (relevance) and expectation (prior) cues (Wyart et al., 2012). The Posner cueing task involved change detection and localization with a single, spatial probabilistic cue (Banerjee et al., 2019; Luo & Maunsell, 2015, 2018; Sagar et al., 2019; Sreenivasan & Sridharan, 2019). This novel conjunction of the two task designs – along with concurrent high-density EEG recordings – enabled answering each of our key questions including: i) the nature of behavioral effects induced by Posner cueing, ii) the precise mechanisms – attentional versus expectational – that mediate these effects, and iii) specific neural signatures – SSVEPs versus alpha oscillations – that underpin these effects. In addition, with a novel deep convolutional neural network, we provide definitive evidence for a match between neural representations of attention and Posner cueing.

**Figure 1.**
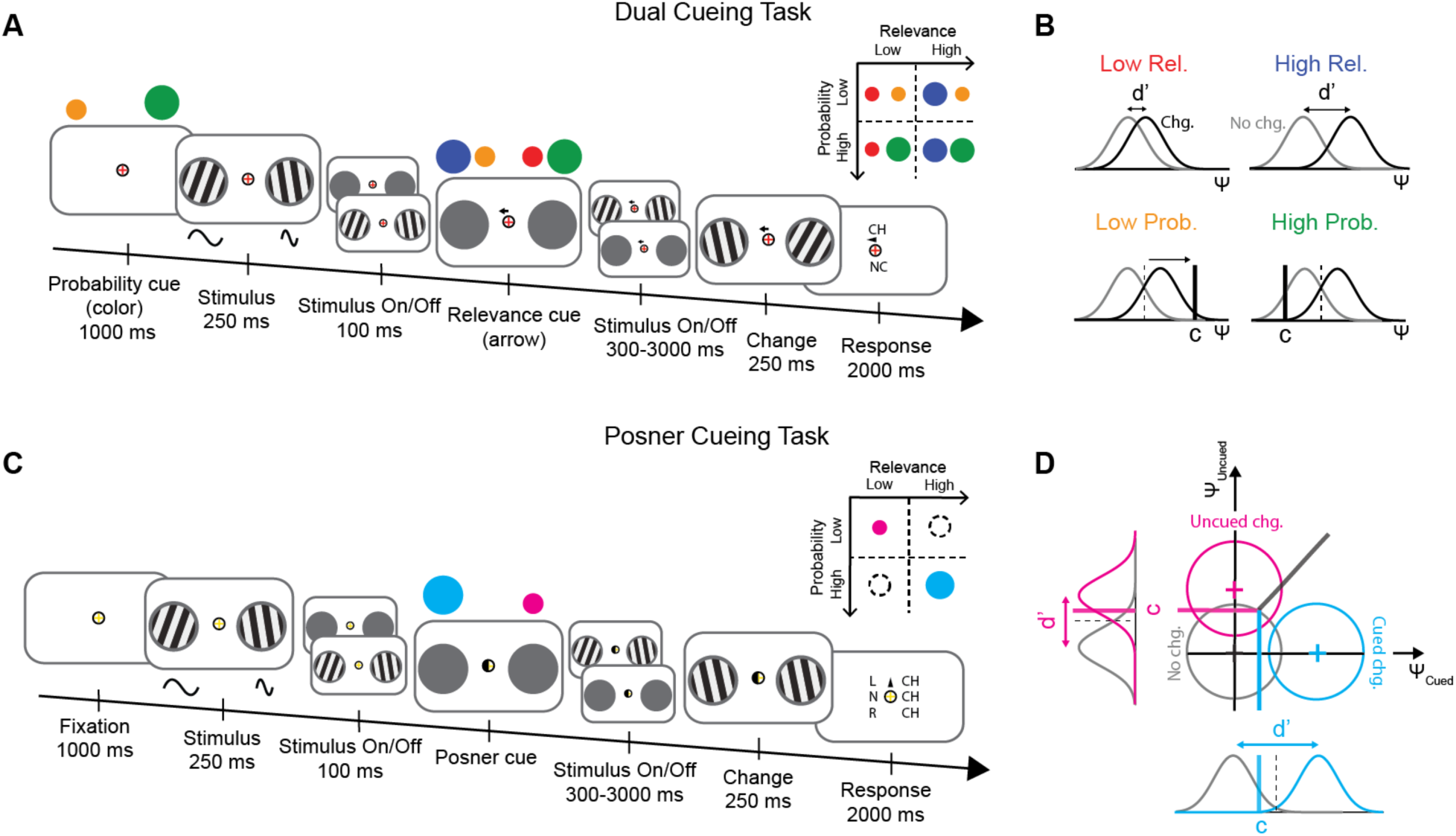
Distinguishing attention from expectation effects in Posner cueing tasks. A. Dual cueing task. Following probability cue (fixation cross color) presentation (1000 ms) two pedestal-mounted Gabor gratings appeared, flickering at distinct frequencies (sine waves, below), one in each visual field. Gratings appeared (250 ms) and disappeared (50-300 ms), alternately (Methods). 350 ms after initial stimulus onset, the relevance cue appeared (central arrow). Following a variable delay (300-3000 ms), either one, both or none of the gratings changed in orientation. Participants reported whether a change had occurred (CH) or not (NC) at the response probed location (arrowhead), based on a trial-specific response mapping. Colored circles: see inset. (Inset) Cue combinations: Low and high event probability (rows, orange and green circles, respectively) and relevance (columns, red and blue, respectively). All four cue combinations were feasible, and equally probable. B. One-dimensional signal detection models to estimate sensitivity and criterion in the dual cueing task. Hypothesis: relevance cueing (top row) and probability cueing (bottom row) dissociably modulate sensitivity (d’, horizontal arrow) and criteria (c, solid vertical line), respectively. Gray and black Gaussians: decision variable distributions for no-change and change events, respectively. C. Posner cueing task. Same as in panel A, except that in each trial, at most only one grating changed in orientation. The Posner cue (shaded semicircle) indicated the more likely change location. Participants localized the change as left (L-CH), right (R-CH) or no change (NC), based on a trial-specific response mapping. (Inset) Cue combinations. Same as in panel A, inset, except that only two cue combinations were feasible. Dotted circles: Infeasible cue combinations. D. A multi-dimensional signal detection model to estimate sensitivity and criterion in the Posner cueing task. Evidence for change at the two locations is represented along orthogonal dimensions (Ψ_cued_, Ψ_uncued_). Gray, magenta and cyan Gaussians: decision variable distributions for no-change, uncued change or cued change events, respectively. Y-shaped decision surface: corresponding decision zones.

## Results

### Hybrid task design to quantify attentional and expectational effects of Posner cueing

To investigate whether endogenous Posner cueing engaged attention or expectation mechanisms, or both, we tested a set of n=21 participants on two, parallel tasks: i) a change detection (Yes/No) task with both relevance and probability cues (“dual cueing task” Wyart et al., 2012) (Fig. 1A), and a ii) a 3-alternative change detection and localization task, with a spatial probabilistic cue (“Posner cueing task”) (Banerjee et al., 2019; Sagar et al., 2019; Sreenivasan & Sridharan, 2019) (Fig. 1B).

In the dual cueing task, participants were presented with two grating stimuli one in each visual hemifield (Fig. 1A). Following an exponential delay (300-3000 ms), the gratings disappeared. Upon reappearance either one, both, or none of the gratings had changed in orientation. Following this, one of the two hemifields was probed for response, and participants reported whether the stimulus on the probed side had changed in orientation or not with a Yes/No response. Attention and expectation were manipulated orthogonally, with a task design closely similar to that of Wyart et al (Wyart et al., 2012). Attention was manipulated with a relevance cue (central, directed arrow; Fig. 1A, fourth panel from left) that indicated the location most likely to be probed for response (80% validity) whereas expectation was manipulated with a probability cue (fixation cross color; Fig. 1A) that indicated the most likely location of change (80% validity). Relevance and probability cueing were rendered independent by employing an orthogonalized pseudorandom sequence for each cue type (Methods); this ensured that the probability cue did not provide any apriori spatial information about the location to be probed on each trial. For this task, sensitivity and criterion were estimated with a conventional, one-dimensional signal detection theory (SDT) model (Green & Swets, 1974) (Methods section on *Psychophysical Parameter Estimation: Dual Cueing Task*) (Fig. 1B, SI Fig. S1A).

Our task differed from, and improved upon, the previous task design of Wyart et al (Wyart et al., 2012), in the following key ways: i) to avoid temporal anticipation of the target, the cue-target (change event) interval was exponentially distributed, rather than constant (Methods section on *Task Description*); ii) to induce sustained attention, both sets of gratings disappeared (50-300 ms, uniform random intervals) and reappeared (250 ms, fixed interval), in a coordinated manner, throughout the trial, mimicking earlier designs; iii) to avoid motor preparation effects of probability cueing, Yes/No responses were mapped randomly to two distinct response keys on each trial; the mapping was revealed 500 ms after probe onset (Fig. 1A last panel; Methods section on *Controlling for Motor Bias*); and iv) to evoke steady-state visually evoked potentials (SSVEPs), each grating was tagged with distinct flicker frequencies (see next section).

The Posner cueing task (Fig. 1C) was nearly identical with the dual cueing task, except that it involved both change detection and localization. On each trial, either one or none (but never both) gratings changed in orientation. Participants localized the change to the left or the right grating, if they perceived a change, or responded “no change” otherwise, with one of 3 button-press responses (buttons randomly mapped to events, see Methods section on *Task Description*). Importantly, in this case, the Posner cue (filled semi-circle; Fig. 1C, fourth panel from left) indicated the most likely location of change (80% validity). Such spatial probabilistic cueing is conventional in “standard attention tasks” and the cue served, therefore, as both a probability and a relevance cue. In this case, we refer to the Posner cued location, and the location in the opposite hemifields, as simply the “cued” and “uncued” locations, respectively. For this task, sensitivity and criterion were estimated with a multi-dimensional SDT model, which has been validated extensively for the analysis of such tasks (Banerjee et al., 2019; Cohen & Maunsell, 2011; Mayo & Maunsell, 2016; Sagar et al., 2019; Sreenivasan & Sridharan, 2019; Sridharan et al., 2014, 2017) (Fig. 1D, SI Fig. S1B) (Methods section on *Psychophysical parameter estimation: Posner cueing task*).

To distinguish attention from expectation effects of Posner cueing, we quantitatively compared psychophysical parameter – sensitivity and criterion – modulations by Posner cueing with those induced by relevance and probability cueing in the dual cueing task.

### Posner cueing mimics attention, not expectation, in its behavioral effects

First, we tested if relevance and probability cueing produced dissociable effects on sensitivity and criterion in the dual cueing task: based on past literature (Wyart et al., 2012) we expected relevance cueing to alter sensitivity and probability cueing to alter criterion.

In line with literature, relevance cueing reliably modulated sensitivity: median sensitivity was significantly higher at the high relevance, as compared to the low relevance, location (p<0.001, Wilcoxon sign rank test; BF>10^2^) (Fig. 2A, top). Moreover, sensitivity for the low relevance location was significantly greater than zero (p<0.001, signed rank test; d’ _low-rel._ = 0.293 ± 0.07, mean ± s.e.m., BF=73.69), indicating that participants did not simply ignore the low relevance location. On the other hand, criterion was unaffected by relevance cueing (p=0.259; BF=0.47) (Fig. 2A, bottom). Conversely, probability cueing reliably modulated criterion: median criterion was significantly lower at the high probability, as compared to the low probability, location (p<0.001; BF>10^2^) (Fig. 2B, bottom). In this case, sensitivity was unaffected by probability cueing (p=0.543; BF=0.25) (Fig. 2B, top).

**Figure 2.**
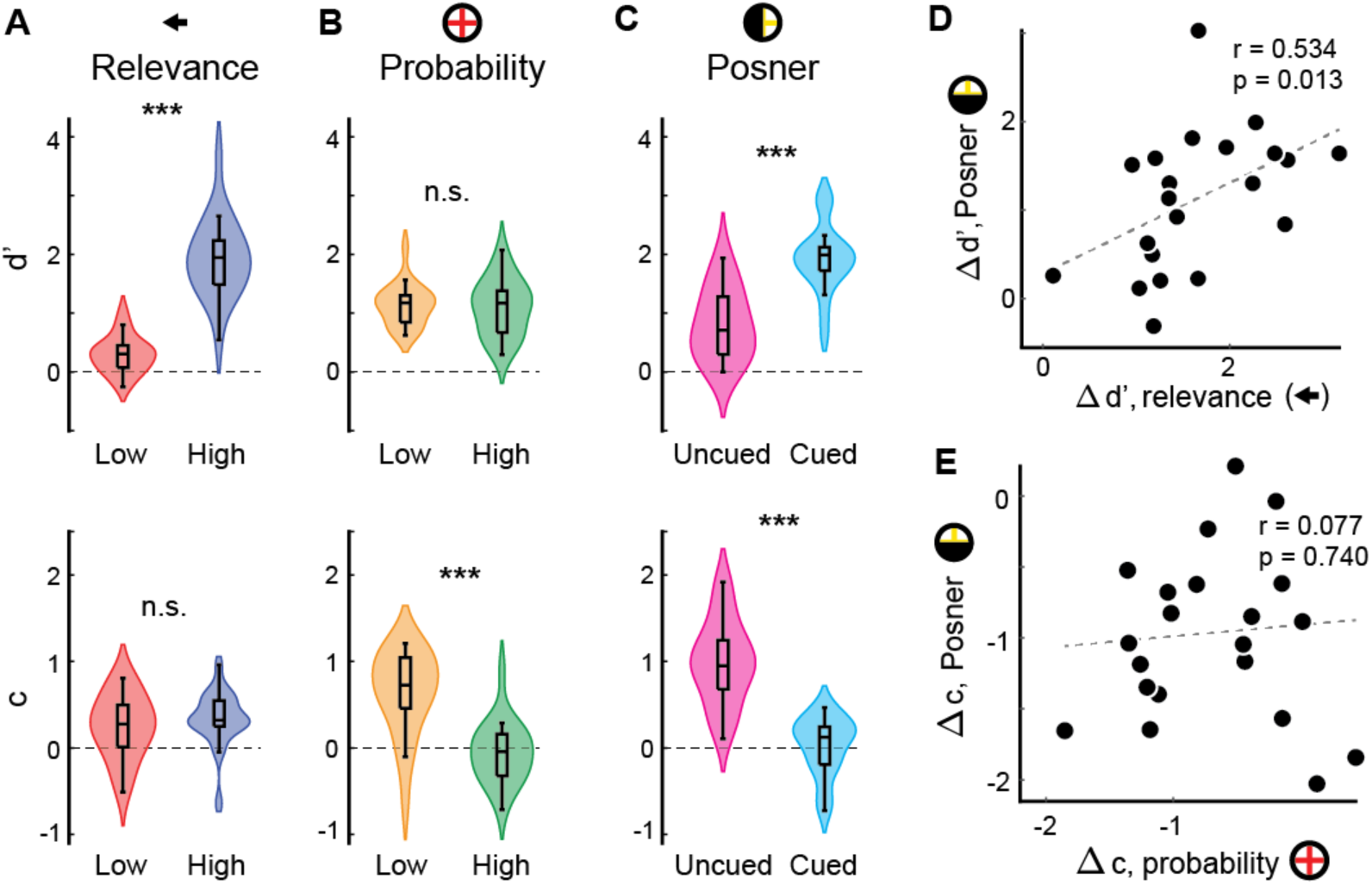
Behavioral effects of Posner cueing reflect relevance, not probability, cueing. A. (Top) Effect of relevance cueing on sensitivity (d’). Red and blue: Low (left) and high (right) relevance conditions, respectively. Violins: data distribution (kernel density estimates using Gaussian kernels) across participants. Lower and upper box edges: lower and upper quartiles, respectively. Mid-line: median. Whiskers: Data range (non-outliers). *p<0.05, **p<0.01, ***p<0.001, n.s.: not significant. (Bottom). Same as in the top panel but showing the effect of relevance cueing on criterion (c). Other conventions are the same as in the top panel. B. Same as in panel A, but showing the effect of probability cueing on sensitivity (top) and criterion (bottom). Orange and green: Low (left) and high (right) probability conditions, respectively. Other conventions are the same as in panel A. C. Same as in panel A, but showing the effect of Posner cueing on sensitivity (top) and criterion (bottom). Magenta and cyan: Uncued (left) and cued (right) conditions, respectively. Other conventions are the same as in panel A. D. Relationship between sensitivity effects of relevance and Posner cueing. x- and y-axes: d’ modulation by relevance cueing (high relevance – low relevance) and Posner cueing (cued – uncued), respectively. Data points: individual participants. r and p value: Percentage bend correlation coefficient and significance level, respectively. Dashed line: best fit. E. Same as in D, but showing the relationship between criterion effects of probability and Posner cueing (x-axis) and modulations in criteria by Posner cueing (y-axis). Other conventions are the same as in panel D.

Although the relevance cue and probability cue were orthogonalized, the relevance cue always followed the probability cue in our task (Fig. 1A), following the task design of Wyart et al (2012). We tested whether this design produced a carryover effect of the (more recent) relevance cue on the probability cue, or vice versa. For this, first, we performed a 2-way ANOVA with relevance level (high, low) and probability level (high, low) as factors. While we found a robust main effect of relevance on d’ (F(1, 20)=109.84, p<0.001) but no main effect of probability (F(1,20)=0.2, p=0.655), we also observed no interaction effect between relevance and probability (F(1,20)=0.47, p=0.501). Similarly, we found a robust main effect of probability on criterion (F(1,20)=31.66, p<0.001) but no main effect of relevance (F(1,20)=1.67, p=0.211); a weak interaction effect between relevance and probability cueing (F(1,20)=4.95, p=0.038), did not survive a Tukey HSD test for multiple comparisons (p>0.1, for all pairwise post hoc comparisons). Next, we also tested whether the congruency of the relevance and probability cues affected behavior. Specifically, we tested whether sensitivity effects of relevance cueing or criterion effects of probability cueing depended on congruence with the other cue type. In all cases, we observed no effects of cue congruence on behavioral effects (d’ modulation by relevance cueing: congruent = 0.802 ± 0.091, incongruent = 0.839 ± 0.086, p=0.543, BF=0.249; c modulation by probability cueing: congruent = 0.031 ± 0.063, incongruent = 0.398 ± 0.080, p=0.259, BF=0.471). Taken together, these results indicate that cue-orthogonalization achieved its desired effect and produced no evident interaction between the cue types.

Moreover, the modulation of sensitivity with relevance cueing (d’_high-rel._−d’_low-rel._) did not correlate with the modulation of criterion with probability cueing (c_high-prob._−c_low-prob._) across participants (r=-0.320, p=0.157, BF=0.30) (SI Fig. S1C). In sum, relevance and probability cueing produced dissociable modulations of sensitivity and criterion, respectively, suggesting that attention and expectation produced independent behavioral effects in this dual cueing task.

By contrast, Posner cueing significantly modulated both sensitivity (p<0.001; BF>10^2^) (Fig. 2C, top) and criterion (p<0.001: BF>10^2^) (Fig. 2C, bottom): median sensitivity was significantly higher, and criterion was lower, at the Posner cued location, as compared to the uncued location. In the Posner cueing task, the location of higher signal probability is also the most relevant location for attention. It is possible, therefore, that behavioral modulations of the psychophysical parameters in the Posner cueing task reflect the effects of attention, expectation, or a mixture of both. Specifically, it is possible that the effects of Posner cueing on sensitivity reflect attentional mechanisms, whereas those on criterion reflect expectation mechanisms. Because each of our participants performed both the Posner cueing and the dual cueing tasks, we tested this hypothesis by analyzing sensitivity and criterion modulations across the two task types.

Sensitivity modulation by relevance cueing in the dual cueing task (d’_high-rel._−d’_low-rel._) was significantly correlated with that by Posner cueing (d’_cued_–d’_uncued_), across participants (r=0.534, p=0.013, BF=3.64) (Fig. 2D). On the other hand, surprisingly, criterion modulation by probability cueing (c_high prob._−c_low prob._) was not correlated with that by Posner cueing (c_cued_– c_uncued_) (r=0.077, p=0.740, BF=0.18) (Fig. 2E). Analysis with the Pearson and Filon’s z statistic (Diedenhofen & Musch, 2015) revealed that sensitivity modulations were more strongly correlated across tasks than criterion modulations (p=0.040). Moreover, criterion modulation by Posner cueing was not correlated with either d’ modulation by Posner cueing (r=-0.160, p=0.489, BF=0.211) nor with d’ modulation by relevance cueing in the dual cueing task (r=-0.135, p=0.559, BF=0.184). In other words, although sensitivity modulations in the Posner cueing task resembled those induced by relevance cueing, criterion modulations in the Posner cueing task did not resemble those induced by probability cueing.

In summary, Posner cueing induced attention-like modulations of d’, indicating that spatial probabilistic cueing engaged attention at the location of higher signal probability. By contrast, although Posner cueing also systematically modulated criterion, these were not correlated either with expectation-linked criterion changes by probability cueing or attention-linked d’ changes by relevance cueing. To further explore these links between Posner cueing and attention or expectation processes, we probed the neural underpinnings of these cues with scalp electrophysiology.

### Posner cueing engages established electrophysiological markers of attention

To understand which process Posner cueing engaged, at a neural level, we quantified and compared the electrophysiological (EEG) markers of attention and expectation with those of Posner cueing. These analyses were conducted on n=19/21 participants for whom EEG data were collected (Methods).

First, we examined electrophysiological markers of spatial attention. Previous studies have shown that selective spatial attention induces a localized enhancement of SSVEP power over the cue-contralateral electrodes (Kim et al., 2007; Morgan et al., 1996; Müller et al., 1998), as well as a lateralization of alpha-band (8-12 Hz) oscillation power over the cue-ipsilateral, relative to the cue-contralateral, electrodes (Foxe & Snyder, 2011; Worden et al., 2000). We tested whether each type of cueing would modulate either of these neurophysiological signatures reliably.

Relevance cueing enhanced SSVEP power robustly (Fig. 3A): the increase was significant in a window locked to cue onset (650 to 1650 ms post-cue; p=0.003, Wilcoxon signrank test; BF=11.620), as well as that locked to change epoch onset (-1050 to -50 ms pre-change; p=0.005; BF=11.737) (Fig. 3A, right). By contrast, probability cueing did not modulate SSVEP power significantly in either epoch (post relevance cue: p=0.759; BF=0.135; pre-change: p=0.911; BF=0.093) (Fig. 3B). To ensure probability cueing effects were not diminished by relevance cue onset, we also tested a pre-relevance cue epoch: 0-350 ms after stimulus, but before relevance cue, onset. Probability cueing did not significantly modulate SSVEP power in this epoch also (p=0.905, BF=0.14). Interestingly, Posner cueing also modulated SSVEP power reliably: the increase was robust in both the post-cue (p=0.018, BF=2.05) and pre-change epochs (p=0.005, BF=2.96) epoch (Fig. 3C). In other words, SSVEP power modulation by Posner cueing resembled the modulation observed with relevance cueing, but not probability cueing, in the dual cueing task.

**Figure 3.**
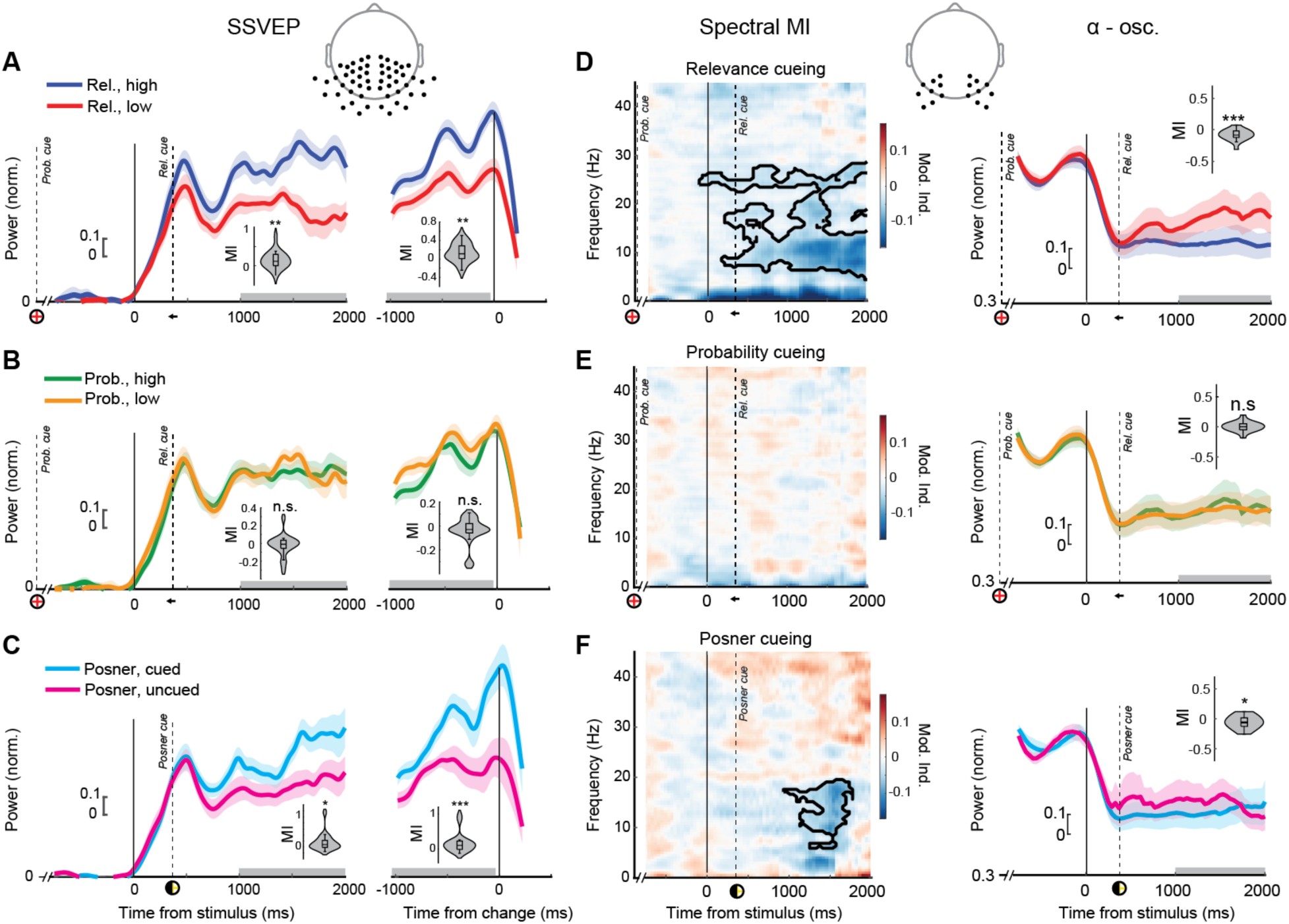
Neural signatures of Posner cueing mimic relevance, not probability, cueing. A. (Left) SSVEP power time courses for high (red) and low (blue) relevance conditions locked to stimulus onset (solid vertical line). x-axis: time from stimulus onset; y-axis: SSVEP power (normalized, Methods). Dashed vertical lines: probability or relevance cue onset. (Inset) SSVEP power modulation index (MI), computed in a 1000 ms time window (gray horizontal shading) following relevance cue onset. Violin plot: conventions are as in Figure 2A. (Right) Same as in the left panel, but locked to change epoch onset (solid vertical line). (Inset) SSVEP power MI, computed in a 1000 ms time window (gray horizontal shading) prior to change epoch onset. *p<0.05, **p<0.01, ***p<0.001, n.s.: not significant. (Inset)Topoplot shows the electrodes used for obtaining SSVEP dimensions using RESS. B. Same as in panel A, but showing SSVEP power time courses for high (green) and low (orange) probability conditions. C. Same as in panel A, but showing SSVEP power time courses for Posner cued (cyan) and uncued (magenta) conditions. (B-C) Other conventions are the same as in panel A. D. (Left) Modulation of broadband spectral power by relevance cueing. x-axis: time from stimulus onset, y-axis: frequency; z-axis: modulation index; warmer colors denote higher values. Black contour: significant spectrotemporal clusters (one-tailed cluster permutation test, p<0.05). E. (Right) Same as in panel A, left, but showing alpha-band power time courses for high (red) and low (blue) relevance conditions locked to stimulus onset. (Inset) Same as in panel A, inset, but showing alpha-band power MI. Other conventions are the same as in panel A. (Inset) Topoplot shows the electrodes used for spectrogram and alpha band power analyses. F. Same as in panel D, but showing broadband spectral power modulation by probability cueing (left) and alpha-band power time courses (right) for high and low probability conditions. G. Same as in panel D, but showing broadband spectral power modulation by Posner cueing (left) and alpha-band power time courses (right) for Posner cued and uncued conditions. (E-F) Other conventions are the same as in panels B-D.

Similarly, relevance cueing produced robust alpha lateralization shortly after cue onset: alpha power increased significantly over cue-ipsilateral, relative to cue-contralateral, occipital electrodes (Fig. 3D, right panel). Spectral power quantification revealed robust lateralization of occipital alpha-band (8-12 Hz) power following the relevance cue onset (650 to 1650 ms post-cue; p<0.001, BF>10^2^) epoch (Fig. 3D, right panel, inset). By contrast, probability cueing produced no significant alpha lateralization during this epoch (p=0.540, BF=0.20) (Fig. 3E, right panel). Posner cueing also produced significant occipital alpha lateralization during the post-cue epoch (p=0.021, Wilcoxon signed rank test; BF=4.06) (Fig. 3F, right panel), again, mimicking the pattern observed with relevance cueing. Similar results were observed with pre-change alpha power also which showed a significant lateralization with relevance and Posner cueing but not by probability cueing. Lastly, following a recent study (Trajkovic et al., 2023), we tested if individual alpha frequency peaks – the frequency corresponding to the peak of the alpha power for each individual for each cueing type – were modulated by relevance cueing, but failed to find clear evidence supporting this hypothesis (p=0.290, BF=0.256); similar results were obtained for Posner cueing also (p=0.190, BF=0.444).

Next, we analyzed the effects of cue congruency on neural signatures. As with the behavior, we did not observe any difference in post-cue SSVEP power modulation by the relevance cue as a result of congruence with the probability cue (SSVEP power modulation: congruent = 0.095 ± 0.069, incongruent = 0.192 ± 0.054, p=0.573, BF=0.624). Similarly, post-cue alpha suppression by the relevance cue did not vary with its congruence with the probability cue (alpha power modulation: congruent = -0.079 ± 0.030, incongruent = -0.095 ± 0.022, p=0.159, BF=0.259).

Finally, we investigated links between these neural signatures of Posner cueing, and its behavioral effects on d’ and c; we also tested whether these links would match with those of either relevance or probability cueing. First, we tested whether the modulation of SSVEP power would predict the modulation of either psychophysical parameter (d’, c) in the dual cueing task. For this, we performed a tercile split of SSVEP power in the pre-change window across trials and estimated d’ and c for the top third (“high”) and bottom third (“low”) trial subsets; this split was performed separately for the subset of trials in which the high relevance or high probability locations were probed (Methods section on *Tercile split analysis*).

d’ for high SSVEP power trials was significantly higher than that for low SSVEP power trials (p=0.017, BF=3.65; one-tailed permutation test) at the high relevance location (Fig. 4A). By contrast, criteria did not vary significantly with SSVEP power at the high relevance (p=0.239, BF=0.64, Fig. 4B). Equivalent results we obtained based tercile split of SSVEP power at the high probability location (p=0.045, BF=5.15 for sensitivity modulations, p=0.232, BF=0.790 for criterion modulations) (SI Fig. S2A,B). A similar analysis with the Posner cueing task revealed strong evidence for d’ variation with SSVEP power (p=0.046, BF=9.49) at the Posner cued location (Fig. 4C). Interestingly, criteria in the Posner cueing task also varied with SSVEP power (p=0.018, Fig. 4D), but the evidence for this effect was anecdotal (BF=1.91). A similar tercile split analysis based on pre-change alpha power revealed no statistically significant modulation of either d’ or criterion, by any type of cueing (SI Fig. S3, see Discussion).

**Figure. 4.**
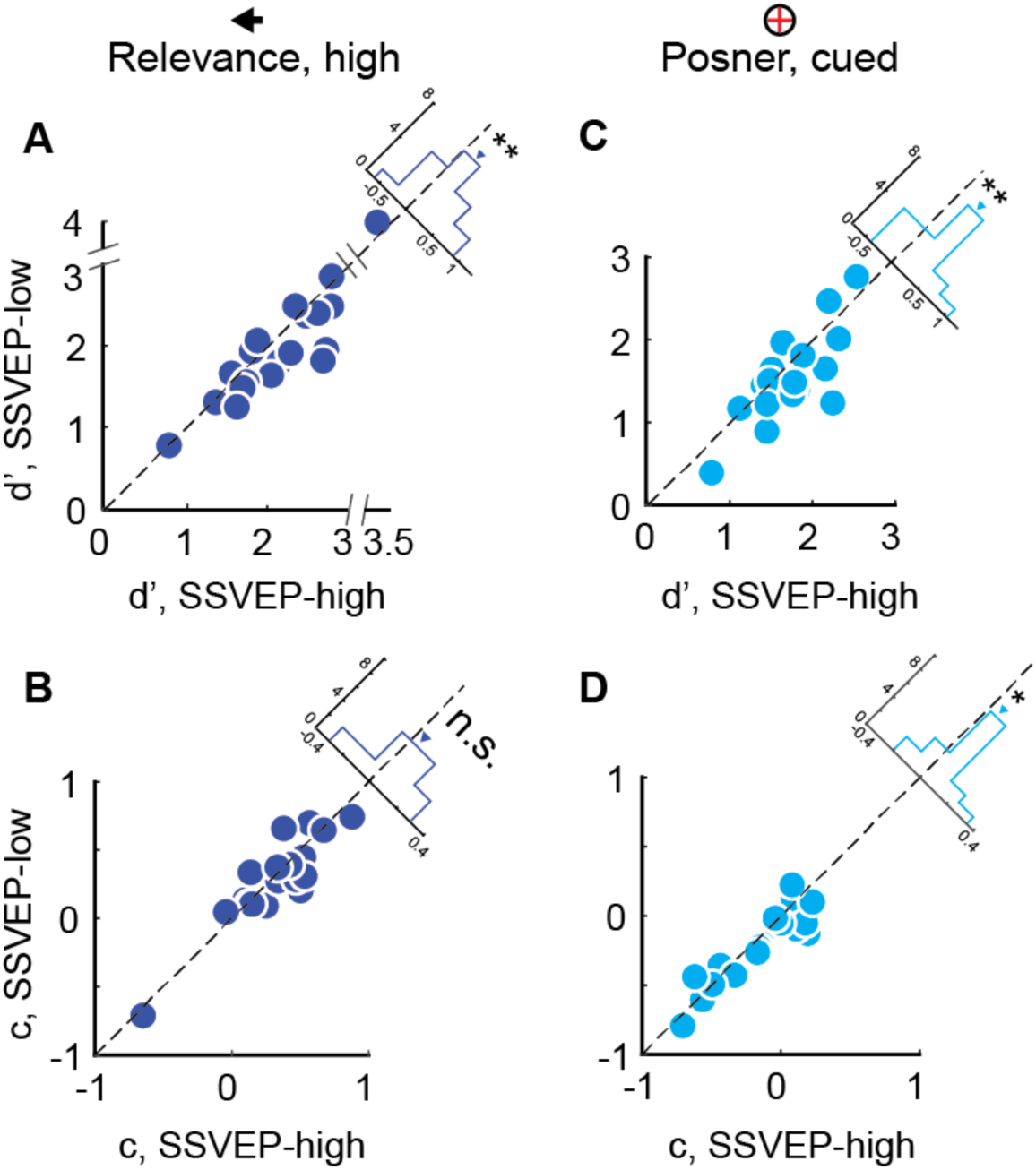
SSVEP power predicts sensitivity variations with relevance and Posner cueing. A. Sensitivities (d’) for trials with highest (top 33%, x-axis) and lowest (bottom 33%, y-axis) terciles of change locked SSVEP power in the dual cueing task (high relevance condition). Data points: participants. Upper right inset: histogram of d’ differences (d’SSVEP-high – d’SSVEP-low). Dashed line: line of equality. *p<0.05, **p<0.01, ***p<0.001, n.s.: not significant. B. Same as in panel A, but showing criteria (c) based on a tercile split of SSVEP power in the dual cueing task (high relevance condition). C. Same as in panel A, but showing sensitivities based on a tercile split of SSVEP power in the Posner cueing task (cued condition). D. Same as in panel A, but showing criteria based on a tercile split of SSVEP power in the Posner cueing task (cued condition). (B-D) Other conventions are the same as in panel A.

Next, we examined electrophysiological markers of spatial expectation; unlike spatial attention there is considerably less consensus on EEG markers of spatial expectation (Doricchi et al., 2010; Tarasi et al., 2022; van Ede et al., 2020; Zuanazzi & Noppeney, 2019). Recent work suggests that increase in pre-stimulus (anticipatory) fronto-parietal theta connectivity or a decrease in pre-stimulus alpha power (Tarasi et al., 2022; van Ede et al., 2020) in the occipital electrodes may index higher signal expectation, but the latter marker has not been consistently replicated (Zhou et al., 2021). In addition, a phase opposition index in the alpha band, which measures the phase locking of band-limited event traces across trials (Sherman et al., 2016), has been found to differentially predict “Yes” and “No” responses. Another recent study (Di Gregorio et al., 2023) observed modulations in fronto-parietal alpha connectivity as an index of post stimulus processing. We tested for the modulation of these neural markers – fronto-parietal theta connectivity, fronto-parietal alpha connectivity, occipital alpha power modulation and alpha phase opposition index – by all three types of cueing (see Methods, section on *Neural markers of spatial expectation*). We examined these signatures in a pre-change (-1050 to -50 ms before change) epoch rather than the pre-stimulus epoch because orientation change was the relevant event of interest for our tasks. These previously reported neural signatures of spatial expectation were, by and large, not evoked by probability cueing in the dual cueing task, nor were they reliably evoked by Posner cueing (Supplementary Results section on *Neural markers of spatial expectation*, SI Fig. S4). We address this null result in the Discussion, and test alternative, data-driven approaches, in the next section.

### Deep learning definitively maps neural signatures of Posner cueing to those of attention

We found clear and consistent evidence showing that behavioral and neural signatures of Posner cueing matched those of relevance (spatial attentional) cueing. Yet, despite clear effects of spatial expectation on behavior, we failed to replicate previously reported electrophysiological markers of expectation (SI Fig. S4). A potential reason is that spatial expectation signatures have been investigated only in a handful of recent studies, without clear consensus emerging (Doricchi et al., 2010; Sherman et al., 2016; Tarasi et al., 2022; van Ede et al., 2020; Zuanazzi & Noppeney, 2019). Nevertheless, it is also possible that some aspect of our own task design precluded expectation signatures from reliably manifesting in the EEG recordings. In this case, it would be challenging to identify neural mechanisms of Posner cueing with attentional mechanisms alone.

To explore this latter possibility, we, adopted two data-driven approaches that did not rely on finding well-documented neural signatures of attention or expectation. First, we employed Representational Similarity Analyses (RSA) (Sheahan et al., 2021; Walther et al., 2016). Specifically, we asked whether RSA could capture expectation signatures, as distinct from attention signatures, in the dual cueing task. Briefly, a neural representational dissimilarity matrix (RDM) is computed to quantify the neural dissimilarity between every pair of conditions across trials; the *β* coefficients obtained by regressing this neural RDM at each timepoint against condition-based “model” RDMs quantify how the evidence for each condition evolves with time (Methods section on *Neural representations of relevance, probability and Posner cueing*). Here, we employed model RDMs for relevance cueing (relevance RDM) and probability cueing (probability RDM), as well as the initial stimulus orientation (visual RDM) (Fig. 5A). The RSA analysis revealed clear evidence for dissociable neural representations of relevance and probability cueing: Evidence for attention signatures peaked following relevance cue onset (Fig. 5B, blue trace) whereas evidence for expectation signatures peaked earlier during the trial, shortly after probability cue onset (Fig. 5B, green trace). In other words, EEG recordings carried clear evidence for dissociable attention- and expectation-linked neural representations during distinct trial epochs.

**Figure 5.**
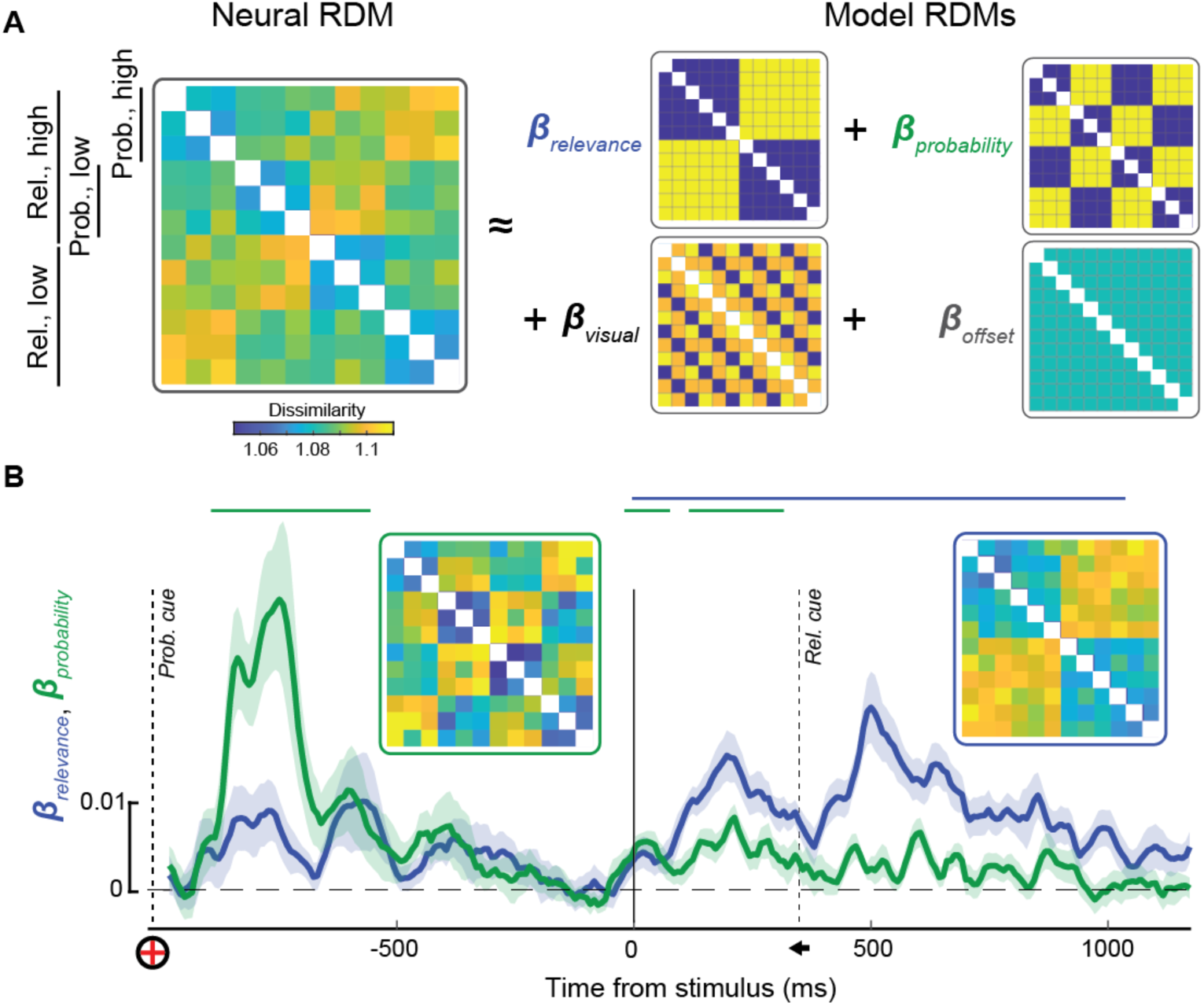
Distinct neural underpinnings of relevance and probability cueing. A. (Left) Neural representational dissimilarity matrix (RDM; average across participants and time) showing dissimilarity (Methods) between each pair of conditions (z-axis). Warmer conditions denote greater dissimilarity; white: self-pairs not included in the analysis. The rows and columns of the neural RDM are ordered and grouped by visual (grating orientation) and cue (probability, relevance) conditions. (Right) The regression model showing the explanatory variables (model RDMs) for each condition and the associated regression coefficients (β_relevance_, β_probability_, and β_visual_). β_offset_: intercept. B. Time course of the regression coefficients for the relevance (blue) and probability (green) model RDMs ((β_relevance_ and β_probability_, respectively). x-axis: time from stimulus onset. Solid vertical line: stimulus onset. Dashed vertical lines: probability or relevance cue onsets. Horizontal lines on top: significant temporal clusters for the probability (blue) and relevance (green) regression coefficients (cluster-permutation tests, p<0.05). Inset RDMs (atop traces): Neural RDMs averaged in a 500 ms window following probability cue onset ([250, 750] ms after cue onset, green outline, left inset), and in a 500 ms window following relevance cue onset (blue outline, right inset).

Second, we developed deep convolutional neural network (CNN) architectures to decode each type of cue from EEG recordings (Methods, Fig. 6A). Briefly, we trained independent 7-layer deep CNN models that were provided with single-trial 93 × 49 × 128 (time × frequency × electrodes) spectrotemporal tensors as input (Fig. 6A, first panel) and predicted, as output, trial labels for the relevance and probability cue locations (left or right) (Fig. 6A, last (blue) panel); we call these the Attention-CNN and the Expectation-CNN, respectively. A novel feature of our model’s architecture was the inclusion of 3 embedding layers: i) an “electrode embedding” (Fig. 6A, ‘elec emb.’) that accounts for variations in electrode positions among participants, ii) a “time-frequency embedding” (Fig. 6A, ‘t x f emb.’) that accounts for variations in spectral peaks and event latencies among participants, and iii) a “participant embedding” (Fig. 6A, yellow box) that accounts for other idiosyncratic sources of behavioral variation (e.g., alertness, motivation) among participants. The model was trained by pooling data across all participants: while the deep CNNs modeled participant-generic relationships between EEG features and cue locations, the embeddings accounted for participant-specific effects. Data from 16,843 trials were included in this analysis, and predictions evaluated with 5-fold cross-validation (Methods, section on *Neural representations of relevance, probability and Posner cueing*) (SI Table S1).

**Figure 6.**
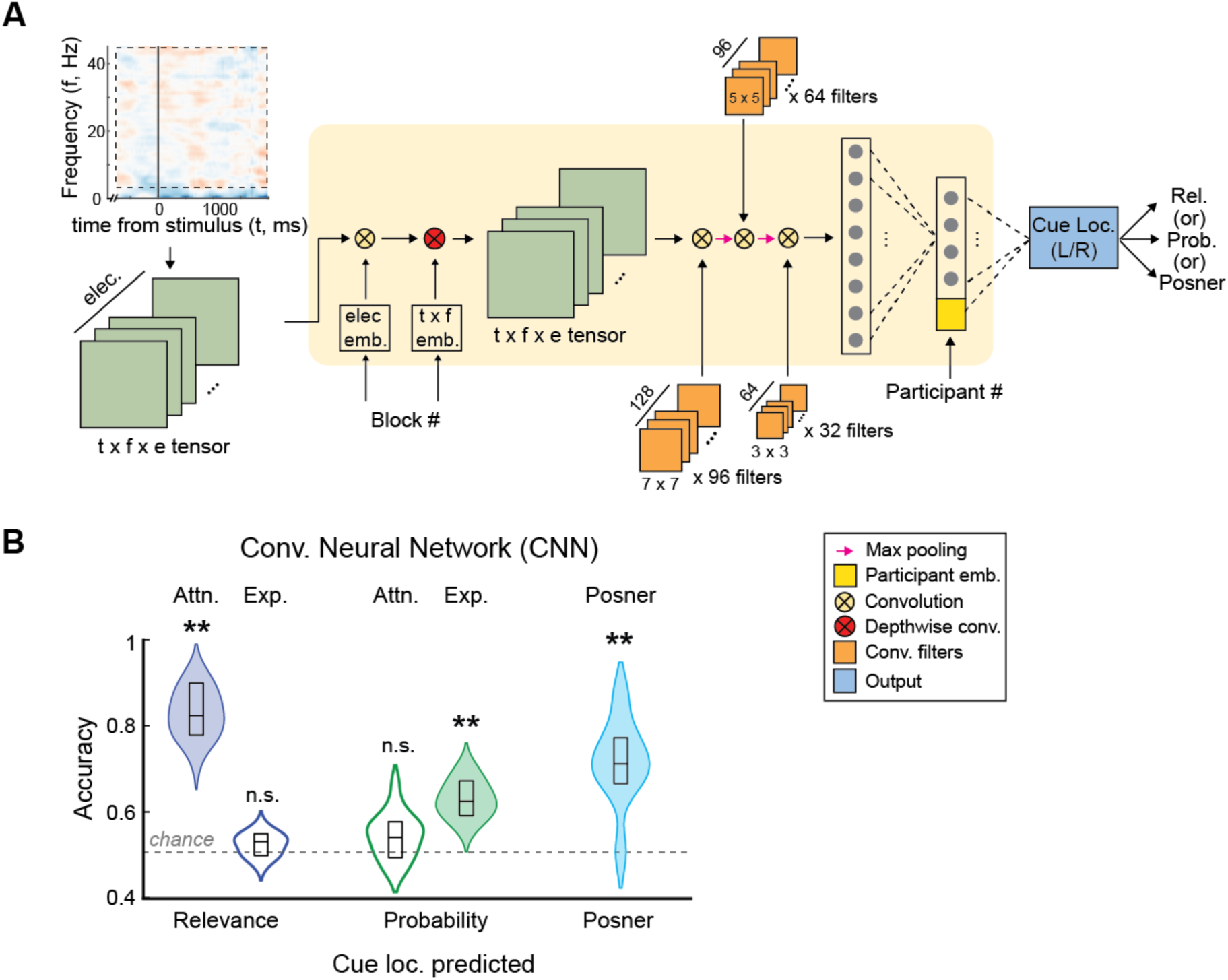
Deep convolutional neural network confirms dissociable cueing representations. A. Architecture of the deep-convolutional neural network (CNN) model. The model receives as input 3D spectrogram tensors (time x frequency x electrodes; leftmost) and is trained to output the predicted cue location (left/right; rightmost). Three independent models were trained for the three types of cues: the Attention-CNN (relevance cues), Expectation-CNN (probability cues) and Posner-CNN (Posner cues), respectively. Yellow box: core architecture with “electrode” and “time-frequency” embeddings (“elec emb.”, “txf emb.”), three convolutional layers with max-pooling (yellow X’s with pink arrows), and two dense layers (tall rectangles). The output of the penultimate layer is concatenated with “participant”-specific embeddings (yellow box in dense layer). The final linear layer predicts the respective cued location (“cue loc. (L/R)”) (see Methods for further details of architecture and training). B. Prediction accuracies of cue locations. (Left) Accuracy for predicting the relevance cue location (blue) by the Attention-CNN (left, filled) or the Expectation-CNN (right, open). (Middle) Accuracy for predicting the probability cue location (green) by the Attention-CNN (left, open) or the Expectation-CNN (right, filled). (Right) Accuracy for predicting the Posner cue location by the Posner-CNN (cyan, filled). Violins: accuracy distribution (kernel density estimates with Gaussian kernels) across participants. Lower and upper box edges: lower and upper 99% confidence intervals, respectively. Mid-line: median. Dashed line: Chance accuracy (50%). *p<0.05, **p<0.01, ***p<0.001, n.s.: not significant.

The Attention-CNN model decoded relevance cue locations significantly above chance (accuracy: median, [99% CI] = 82.35, [77.84, 89.85]%, n=19 participants; 25 test sets with 5 seeds x 5 folds per seed) (Fig. 6B, leftmost). Similarly, the Expectation-CNN decoded probability cue locations with 61.91 [58.65, 66.63]% accuracy, again, significantly above chance (Fig. 6B, penultimate violin plot). Next, to test if the decoding of relevance and probability labels were driven by overlapping or distinct features, we probed cross-decoding performance across these models. Both Attention-CNN and the Expectation-CNN were not able to predict the other’s labels above chance level accuracy (54.32 [49.58, 57.9]%, for Attention-CNN predicting probability cue locations; Fig. 6B, second violin plot from left and 52.47 [49.23, 54.25]%, for Expectation-CNN predicting relevance cue locations; Fig. 6B, third violin plot from left). In other words, each CNN decoder predicted the likely spatial locations of high attention and high event expectation, respectively, with above chance accuracies. Yet, neither was able to decode the other’s labels reliably, suggesting that the CNN models relied on largely distinct, non-overlapping features to predict their respective cue locations. These results confirm the independent neural representations of attention and expectation identified with the RSA analysis.

Next, we trained a third, independent deep CNN model to predict Posner cue locations, again from single trial spectro-temporal tensors; data from 7,200 trials were included in this analysis (Methods). The “Posner-CNN” decoded the Posner cued location with 70.39 [65.87, 76.43]% accuracy, significantly above chance, indicating robust representation of Posner cue-related information in the EEG traces (Fig 6B, last violin plot).

Finally, we tested whether neural representations of Posner cueing matched those of relevance cueing or probability cueing. For this, we tested whether the features employed by the Posner-CNN for label prediction, would correspond more with those of the Attention-CNN or with those of the Expectation-CNN. Briefly, we computed “saliency maps” (Simonyan et al., 2013) which enable visualizing which features in the spectro-temporal tensor were helpful for discriminating among the labelled classes (Methods, section on *Neural representations of relevance, probability and Posner cueing, Part (ii*)*: Saliency Maps*). Saliency maps for both the Attention-CNN and Expectation-CNN identified, among other features, prominent weights in a low-frequency band in the alpha/low-beta range (10-20 Hz), as well as a high-frequency band in the high-beta/gamma range (30-40 Hz) (Figs. 7A-B). A key difference was that, temporally, the highest weights for the Attention-CNN occurred following relevance cue onset (Fig. 7A), whereas for the Expectation-CNN the highest weights occurred following probability cue onset (Fig. 7B). These results are in line with results from the previous RSA analysis (Fig. 5B, blue trace for *β*_relevance_, following relevance cue onset and *β*_probability_, green trace, following probability cue onset).

**Figure 7.**
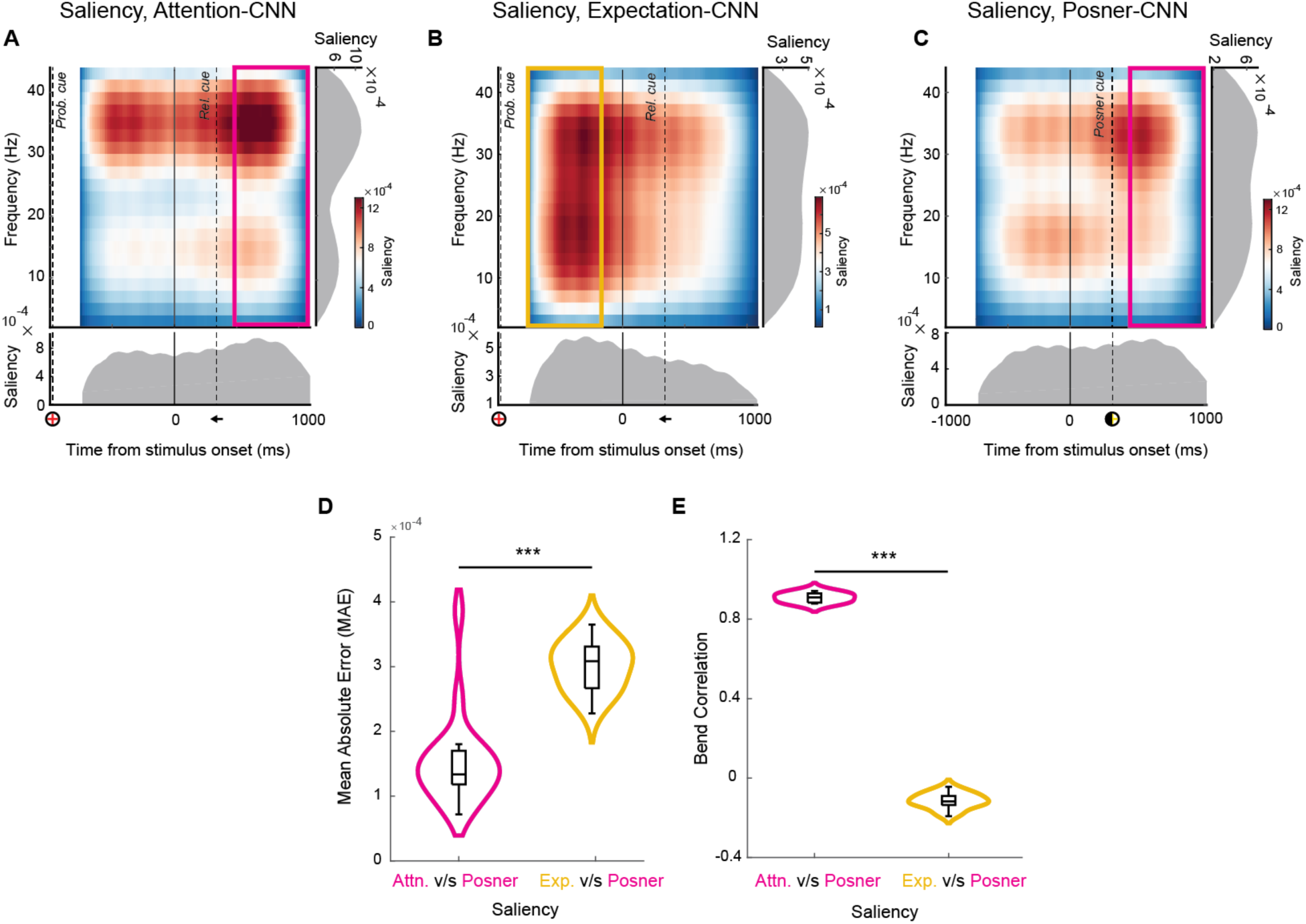
Posner-CNN saliency maps match Attention-CNN maps. A. Saliency map for the Attention-CNN showing time-frequency combinations critical for predicting relevance cue labels. z-axis: salience; warmer colors indicate higher salience, x-axis: time from stimulus onset, y-Axis: frequency. Solid vertical line: stimulus onset; dashed vertical lines: probability or relevance cue onsets. Magenta color box: time-frequency windows used for computing mean absolute error and correlations with the Posner saliency map. Insets along each axis: Time (below) and frequency (right) marginal saliency distributions B. Same as in panel A, but showing saliency maps for the Expectation-CNN. Yellow color box represents the saliency used for computing mean absolute error and correlations with the Posner post-cue saliency map. C. Same as in panel A, but showing saliency maps for the Posner-CNN. Magenta box represents the window used for comparison with the attention post-relevance cue and expectation post-probability cue saliency maps. (B-C) Other conventions are the same as in panel A. D. Mean Absolute Error (y-axis) between salience values of the Attention-CNN and Posner-CNN in post-relevance cue windows (magenta color boxes in panels A and C) and salience balues of the Expectation-CNN and Posner-CNN (yellow box in B and magenta box in C). ***p < 0.001. E. Same as in panel D but showing a bend correlation measure. ***p < 0.001

The saliency map for the Posner-CNN resembled the Attention-CNN saliency map more than the Expectation-CNN map, visually (Fig. 7A-C). To quantify this similarity, we computed the mean absolute error (MAE) and correlations between saliency maps. Because we observed signatures of expectation (attention) closely following the probability (relevance) cue onset (Fig. 5B), we compared the saliency maps of the Expectation-CNN in a post-probability cue window (yellow box, Fig. 7B) or the Attention-CNN in a post-relevance cue window (magenta box, Fig. 7A) with the Posner-CNN in a post-Posner cue window (magenta box, Fig. 7C). MAE was lower (p<0.001), and correlation between maps was higher (p<0.001) between Attention-CNN and Posner-CNN saliency maps than between the Expectation-CNN and Posner-CNN saliency maps (Fig. 7D,E). These results suggest that neural representations recruited by Posner cueing more closely resembled those of relevance cueing than those of probability cueing.

In summary, relevance and probability cueing produced dissociable effects on behavior, with the former modulating sensitivity and the latter modulating bias in our task. Posner cueing modulated both psychophysical parameters. Only sensitivity modulations produced by Posner cueing were correlated with those produced by relevance cueing. SSVEP power and alpha lateralization – established neural markers of attention – were modulated by both relevance and Posner cueing but not by probability cueing. Representation similarity analyses revealed dissociable signatures of attention and expectation, a result confirmed by cue-label prediction with deep convolutional neural networks. Additionally, comparing saliency maps of the deep networks showed that Posner cueing representations more closely resembled those of relevance cueing than those of probability cueing. Taken together, these results demonstrate that Posner cueing engages attention, not expectation, mechanisms.

## Discussion

Although various cueing protocols are effective for guiding selective spatial attention (Carrasco, 2011), spatial probabilistic “Posner” cueing is among the most common in studies investigating the neurophysiological basis of this phenomenon (Ciaramitaro et al., 2001; Cohen & Maunsell, 2009; Cook & Maunsell, 2002; Giordano et al., 2009; Quest et al., 2022; Raffi & Siegel, 2005; Speed & Haider, 2021; Wang & Krauzlis, 2018). In the Posner paradigm, prior knowledge of event occurrence is the only cue guiding the selection of a location for spatial attention. Indeed, seminal studies have argued that “*the subject’s knowledge about where in space a stimulus will occur affects the efficiency of detection*” (Posner et al., 1980), and that, “*with advance knowledge, the mental spotlight can be turned to the correct position before the stimulus is presented*” (Duncan, 1984).

Nonetheless, it has been reasonably argued that the behavioral effects of Posner cueing, particularly in detection tasks, could arise from conflation of attention and expectation (Wyart et al., 2012). In particular, criterion-related effects may arise from a higher signal expectation at the Posner cued location – this being the most likely location of the event of interest (Carrasco et al., 2002; Eckstein et al., 2002; Morgan et al., 1998). In fact, Posner et al (Posner et al., 1980) mention this confound: *“… [subjects] may seek to raise their criterion at the unexpected position and lower it at the expected position. This might have nothing to do with capacity or attentional limitations but would simply be an adaptation to the experimental contingencies.*”

Providing suggestive evidence for the latter hypothesis, Wyart et al (Wyart et al., 2012) designed an elegant binary choice (Yes/No) detection task in which attention and expectation were decoupled using distinct relevance and probability cues, respectively. With a conventional signal detection theory (SDT) model, they observed dissociable effects of these two processes on behavior: relevance cueing modulated SDT sensitivity alone whereas probability cueing modulated SDT criterion alone (Wyart et al., 2012). More specifically, their fine-grained experimental analyses revealed a higher “energy sensitivity” of false positives, while their modeling analysis revealed evidence consistent with a “baseline shift”, rather than a “threshold shift”; both of these findings are consistent with an SDT criterion, but not an SDT sensitivity (signal-to-noise ratio), change. We replicated these seminal findings in our study, while additionally controlling for a potential motor bias confound on criterion effects, by randomizing response mappings on each trial. Taken together with other recent evidence (Tarasi et al., 2022) these studies convincingly demonstrate that when a high signal probability location is rendered task irrelevant, by design, attention is not directed toward this location.

Yet, what if spatial priors about task relevant events provide the only information for guiding spatial attention? Again, this is precisely the scenario in Posner cueing tasks. In such tasks, it is essential to decouple sensitivity and criterion effects; but conventional, one-dimensional signal detection models cannot achieve this decoupling (Green & Swets, 1974; Macmillan & Creelman, 2005). With a recent multidimensional signal detection model (Sridharan et al., 2014, 2017), we show that Posner cueing systematically modulates both sensitivity and criteria, while also producing clear dissociations between them. Posner cueing yielded attention-like enhancements on sensitivity, matching relevance cueing effects (Fig. 2A, C, top). Yet, remarkably, criterion modulations produced by Posner cueing were not correlated with those by probability cueing (Fig. 2D). In other words, criterion modulations by Posner cueing do not appear to arise from expectation mechanisms.

Behavioral modulations of d’ and criterion were accompanied by modulations of established electrophysiological markers of attention. SSVEP power increased with relevance cueing – matching observations in literature (Morgan et al., 1996; Müller et al., 1998) – as well as with Posner cueing. Similarly, alpha-band power lateralization – a widely-reported marker for spatial attention (Foxe & Snyder, 2011; Rihs et al., 2007; Thut et al., 2006) – occurred robustly with both relevance cueing and Posner cueing. Neither of these signatures occurred with probability cueing. Moreover, probing the behavioral correlates of the SSVEP modulations revealed that trials with higher SSVEP power predicted higher sensitivity, both at the relevance cued and at the Posner cued locations. These results are consistent with recent studies which propose SSVEP power as a reliable marker for sensitivity enhancements induced by endogenous spatial attention (Chen & Golomb, 2023; Chinchani et al., 2022).

Interestingly, we observed a link between SSVEP power and criteria also in the Posner cueing task: trials with higher SSVEP power manifested higher criteria (Fig. 4D), suggesting that criteria in Posner cueing tasks are governed by attention-linked processes. Recent work (Banerjee et al., 2019) demonstrated a detailed theoretical analysis showing that criteria in change detection and localization tasks – like the one used in the present study – are governed by “differential risk-curvature”. Briefly, overall accuracy, and payoffs, are more susceptible to criteria at the Posner cued location, as compared to the uncued location, due to a higher risk curvature at the cued location. This yields a spatial choice bias favoring the cued location. Furthermore, event probabilities cannot be controlled independently across locations, by design, in the Posner cueing task; as a consequence, criterion modulation in the Posner cueing task may reflect the operation of a capacity-limited mechanism. These results may explain why SSVEP power levels – a marker of attentional selection – were predictive of criterion modulation by Posner cueing, but not by probability cueing. Nonetheless, consistent with earlier studies, criterion modulations in the Posner cueing task were uncorrelated with sensitivity modulations in the same task (Banerjee et al., 2019; Sagar et al., 2019; Sreenivasan & Sridharan, 2019). These results suggest that despite sharing a common neural marker of spatial attention (SSVEP power), sensitivity and criterion modulations with Posner cueing may originate from distinct control mechanisms in the brain.

Our study provides the first report, to our knowledge, of a null effect of spatial expectation on SSVEP power. In general, there is little consensus regarding neural indices that are reliably modulated by spatial expectation. We sought to replicate three different anecdotal neural markers of expectation, reported previously; yet, we did not observe strong evidence for any of these markers. There could be several reasons for these differences. First, many previous studies investigated neural signatures of feature expectation (Gordon et al., 2019; Kok et al., 2017; Kok, Jehee, et al., 2012; Rungratsameetaweemana et al., 2018), whereas studies of spatial expectation are comparatively rare. Second, many studies report expectation violation signatures following the onset of the event of interest (Kok, Jehee, et al., 2012; Rungratsameetaweemana et al., 2018; Todorovic et al., 2011; Zhou et al., 2020). Our task precluded analyses of these signatures because of the limited temporal window following change epoch onset prior to the response probe. Nevertheless, our results are more in line with Zhou et al (Zhou et al., 2021) who also failed to replicate previously reported neural markers of expectation (but see Tarasi et al, 2022). A third possibility is that the null results we observed reflect a limitation of our data or choice of experimental design.

To test and discount task design limitations, we adopted two state-of-the-art, data driven approaches: representational similarity analysis, and a novel, embedding-aided deep learning model. Both methods provided converging evidence for dissociable neural representations governing attention and expectation in the dual cueing task. In both cases, the emergence of these neural signatures coincided with, and followed, the onset of the respective cue type (Figs. 5B, 6B, 7A-B). Moreover, neither the Attention-CNN nor the Expectation-CNN could predict each other’s cue locations suggesting orthogonal neural representations underlying the two processes. In other words, expectational representations were robustly identified as distinct from attentional representations by two, independent approaches, suggesting that the failure to replicate previous expectation signatures was unlikely to be a consequence of our specific task paradigm. Broadly, these results are in line with earlier work which suggests that preparatory activity signaling expectation and attention are at least partially decoupled (Peñalver et al., 2023), and that each process may rely on distinct modes of information integration during hierarchical perceptual inference (Gordon et al., 2019).

We, then, sought to test which of these neural representations matched those of Posner cueing. Yet, performing a direct cross-decoding analysis was ill-advised because the dual cueing and Posner cueing tasks were performed in different experimental sessions; misalignments in input tensor features yielded poor generalizability across the two task types (see Methods for details). To circumvent this limitation, we compared salient features employed by the Attention and Expectation-CNN model for their respective cue location decoding with that of the Posner-CNN model. This analysis revealed significantly higher similarity between the Posner and relevance cueing feature representations than between those of Posner and probability cueing.

In sum, our study shows that spatial probabilistic, Posner cueing selectively engages attentional processes: behavioral and neural consequences of Posner cueing match those of attention, rather than expectation. Overall, our study resolves a fundamental question about mechanisms governing a ubiquitous attention cueing paradigm and enables interpreting its behavioral and neural effects reported widely in literature.

## Materials and Methods

### Participants and ethics approval

Twenty-one participants (5 female; median age: 24 years; range: 21–31 years) with no known history of neurological disorders and with normal or corrected-to-normal vision participated in the experiments. All participants provided informed, written consent, and all experimental procedures were approved by the Institute Human Ethics Committee at the Indian Institute of Science, Bangalore.

### Data exclusion

Behavioral data from all n=21 participants were analyzed, except for trials excluded due to poor fixation (see Methods section on *Eye-tracking).* EEG data was acquired for n=19/21 participants. Of these, for 2/19 participants 5% and 15% of trials in the dual cueing task, respectively, were irretrievably lost because of trigger failures during EEG acquisition. For one more participant, data from two dual cueing blocks were lost due to data corruption during EEG file conversion. These data were excluded from all EEG analyses.

### Behavioral task design and data acquisition

#### Task description

Each participant performed both versions of cueing tasks: a change detection (Yes/No) task with dual cueing and a change detection and localization task with Posner cueing. Participants were seated in an isolated room, with their head placed on a chin rest, and eyes 60 cm from a contrast-calibrated visual display (24-in. BenQ LCD monitor, X-Rite i1 Spectrophotometer). Stimuli were programmed with Psychtoolbox (version 3.0.11) (Brainard, 1997; Kleiner, 2007; Pelli, 1997) using MATLAB R2017b (Natick, MA). Responses were recorded with an RB-540 response box (Cedrus). Participants were instructed to maintain fixation on a central cross throughout each trial during the experiment.

In the dual cueing task involving change detection, each trial began with the presentation of an encircled fixation cross (0.5° diameter, Fig. 1A). The color of the fixation cross served as the probability cue (see next). 1000ms after fixation onset, two placeholders (radius: 3.027 dva) – with superimposed sinusoidal grating stimuli (radius: 2.5 dva; spatial frequency: 8 cpd) – appeared, one in each hemifield; pedestal and grating centers were +/- 5 dva from the fixation cross along the azimuth. Each placeholder flickered at a different frequency to evoke distinct EEG SSVEP responses (see next section); the grating in each hemifield flickered in phase, and at the same frequency as the corresponding placeholder. The orientations of the gratings were drawn independently of each other from a uniform random distribution (-60 to 60 degrees relative to a vertical axis). 350 ms after the stimulus onset, a central cue (arrowhead, Fig. 1A, fourth panel from left) appeared which pointed either to the left or the right side (relevance cue). After a variable delay (300-3000ms drawn from an exponential distribution), the screen was blanked and upon reappearance both, none or either of the gratings could have changed in orientation (change/no-change event). After a fixed delay (350 ms) a probe appeared on the screen indicating the side that was relevant for response, and after another brief delay (500 ms), the response mapping was revealed (Fig. 1A, last panel). Participants responded by pressing one of two response keys (Up/Down) on the button box to indicate whether they perceived a change on the probed side or not; the mapping between response keys and possible responses (Yes/No) was pseudorandomly changed on each trial (see section on *Controlling for motor bias*). To ensure sustained attention at the relevance cued location throughout the trial the grating stimuli were flashed on and off periodically: stimuli were presented for 250 ms and disappeared for a random duration distributed uniformly between 50-300 ms, then reappeared again for 250 ms and so on, until the change/no-change event occurred. Pedestals were presented throughout the trial. In this dual cueing task, the probability cue indicated the side with a higher probability of change occurrence (80% validity). The probability cue was either red or blue in color; the high probability color (red versus blue) was counterbalanced across participants, who were informed about this mapping beforehand. The relevance cue (central arrowhead) indicated the side more likely to be probed for response (80% validity). Relevance and probability cues were generated using pseudorandom sequences for each cue type, which were orthogonalized *post hoc* with a heuristic approach. This ensured that the relevance and probability cues conveyed independent information and that, statistically, the occurrence of either type of event on one side could not be used to predict the event that occurred on the other side. As a result, the joint probability of a location being both high probability and high relevance (0.25), was determined to be exactly the product of marginal probabilities of it being either high probability (0.5) or high relevance (0.5). Similarly, the other three joint probabilities for each location – i) high relevance and low probability, ii) low relevance and high probability and iii) low relevance and low probability – were all determined to be exactly 0.25, the product of their respective marginal probabilities. Although we closely modelled this dual cueing task along the lines of Wyart et al (Wyart et al., 2012) with a view to replicating their behavioral results, our task differs in multiple, salient respects (see Results section on *Hybrid task design to quantify attentional and expectational effects of Posner cueing*, and Methods section on *Controlling for motor bias)*.

The Posner cueing task (Fig. 1B) followed a virtually identical structure as the dual cueing task, expect for the following key differences: i) during the change/no-change event either one or none (but not both) of the stimuli could have changed in orientation; ii) the Posner cue (filled semi-circle; Fig. 1B, fourth panel from left) indicated the likely location of upcoming change (80% validity), iii) Participants indicated which event had occurred – change in the left versus right hemifield grating, or no change – with one of 3 response keys; the center key was mapped to no-change whereas the Up/Down keys were randomly assigned to left or right change, independently across trials (Fig. 1B, last panel); iv) the fixation cross color was uninformative (yellow) in this task.

#### Controlling for motor bias

Our dual cueing task closely mimics the design of Wyart et al (2012) (Wyart et al., 2012), who reported robust effects of probability cueing on criterion, but not sensitivity. Yet, a particular design choice renders criterion modulations in Wyart et al (2012) susceptible to motoric response bias. In their detection task, participants responded with the index finger of each hand for ‘No’ responses and with the middle finger for ‘Yes’ responses. They responded with their right hand if probed for the right stimulus and vice versa. Colored placeholders indicated the location of higher stimulus probability, at least 2 seconds before stimulus onset. In this design it is possible – and even likely – that participants adopted the strategy of preparing a motor response immediately following placeholder onset, based on differences in spatial priors.

For example, if, on a given trial, the right side was cued as a high probability location, then the most likely response effectors on that trial would be the right middle and left index fingers; these fingers could be primed for a response >1.5 seconds before the actual response epoch. The subsequent appearance of the relevance cue would enable even stronger selection of the effector to be primed for response. For example, a rightward (leftward) cue on the aforementioned trial would overwhelmingly favor the right middle (left index) finger as the most likely effector by a factor of 4x as compared to the least likely effector, and by a factor of 2x as compared to the other effectors; critically participants could take advantage of this information at least 2500 ms before the actual response. To avoid the effect of motor bias, we re-mapped the response keys pseudorandomly on each trial; the mapping was revealed to participant toward the end of the trial (Figs. 1A, B, last panel). While this design permitted robust estimation of choice bias without the confounding effect of motor bias, it precluded reliable estimates of reaction times because of the fixed delay with revealing the response mappings on each trial.

#### Training and testing

The experiment was conducted over a period of three days. On the first day, we staircased and trained the participants. Participants were staircased for 40 blocks (10 trials each) of the change detection (Yes/No) task; the change angle corresponding to ∼70% accuracy was used for subsequent experiments. Next, participants performed two 100-trial training blocks each of the dual cueing and Posner cueing tasks (400 trials, total). During these training blocks, participants received on-screen feedback regarding their accuracy and the correct response.

On the second and third days, participants performed one block each of the Posner cueing and the dual cueing tasks: Because, in the dual cueing task, the relevance cue and probability cue conveyed independent information we needed at least 2x the number of trials in this task, so as to have a comparable number of trials when measuring cueing effects for each of the 3 cue types. Moreover, pilot experiments revealed that the Posner cueing task was easier to perform and provided reliable parameter estimates with fewer trials than the dual cueing task. Therefore, Posner cueing task blocks comprised 200 trials each, whereas the dual cueing task blocks comprised 500 trials each. Thus, overall, participants performed 1400 trials of both tasks over the two days. The order of the Posner and dual cueing tasks was counterbalanced across participants and days. Participants performed these trial blocks in contiguous stretches of 100 trials with brief rest period (∼1-2 min) in between; a longer rest period (∼15 min) was provided between the dual and Posner cueing blocks.

In addition, participants found it challenging to follow information provided by the two cues in the dual cueing task when these were pseudorandomly interleaved across trials. To facilitate tracking cue-related information we blocked the probability cue in mini-blocks of 25 trials each, and the relevance cue (independently) in mini-blocks of 50 trials each. Pilot experiments confirmed that such blocking produced far more robust effects on behavioral parameters than pseudorandomly interleaving the two cue types. To maintain a parallel structure, Posner cues were blocked either in 25 trial or 50 trial mini-blocks, one each among the two Posner cueing task blocks, respectively.

#### Eye-tracking

Participants’ gaze was monitored monocularly with a 1000-Hz infrared eye-tracker (EyeLink 1000-Plus) and the data were analyzed offline. Trials in which the participants’ median gaze deviated by >1° from the fixation along the azimuthal direction during the post-relevance cue epoch were flagged. For 7/21 participants eye-tracking rejection rates were high (>15%), likely because these participants all wore eyeglasses. For these participants, we included all trials in the analyses reported in the main text; repeating the analyses with eye-tracking trial rejection for these participants yielded closely similar results. For the remaining 14/21 participants, average rejection rates were 7.3 ± 1.0%, and trial rejection was performed based on gaze deviation, with the aforementioned criteria.

### Behavioral data analysis

#### Psychophysical parameter estimation: Dual cueing task

We employed conventional, one-dimensional signal detection theory (SDT) (Macmillan & Creelman, 2005) to analyze participants’ responses in the dual cueing task involving change detection (Yes/No response, Fig. 1C). We estimated change detection sensitivity (*d*′) and criterion (*c*) at each location and for each condition – whereas *d′* measures detection ability, *c* quantifies the bias for reporting a change, and is inversely related to it. Standard one-dimensional formulations were used for computing *d′* and *c* based on hit rates (change detected correctly) and false alarm rates (change reported on a no-change trial), using a 2x2 stimulus response contingency table (SI Fig. S1A).

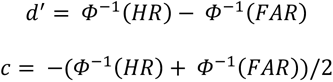

where *Φ*^-1^ is the probit function representing the inverse of the cumulative normal distribution. *d′* and *c* for each location and condition (high/low relevance, or high/low probability) were computed by pooling across trials of the other condition. We also measured the modulation of each psychophysical parameter by each type of cue by taking the difference of the respective parameter across the high and low validity conditions for the respective cue type. For example, Δd’_rel._ = d’_high rel._ – d’_low-rel._

#### Psychophysical parameter estimation: Posner cueing task

We employed a multidimensional SDT model – the m-ADC model(Sridharan et al., 2014, 2017) – to analyze participants’ responses in the Posner cueing task. Conventional, one-dimensional SDT models do not suffice to quantify sensitivity and criterion even in Posner cueing tasks: even the simplest version of this task, with only two potential stimulus locations (cued and uncued), is a multialternative task. For example, in our “2-ADC” task (Fig. 1D) the participant needed to make one of three choices on each trial: change at the cued location, change at the uncued location, or no change. Such tasks cannot be adequately fit with a combination of one-dimensional signal detection models; the reasons are elaborated in Sridharan et al 2014 (p. 3–5) (Sridharan et al., 2014). Here, we recapitulate essential features of the 2-ADC model; a detailed, quantitative description of the more general m-ADC model, for any number of choices, is available in (Sridharan et al., 2014, 2017).

In the 2-ADC task, the participants must detect and localize an orientation change that can occur at either one of the two locations or not at all (no change). From participants’ response proportions for each change event type, we construct a 3x3 stimulus-response contingency table (3 change events x 3 responses). The table comprises five different categories of responses: 1) Hits – change trials in which the participant correctly localized change; 2) Correct Rejections – no change trials which the participants correctly reported; 3) Misses – change trials in which participants reported no change; 4) False Alarms – no change trials in which participants reported a change; 5) Mislocalizations – change trials in which participants localized the change to an incorrect location. To fit the m-ADC model for each participant we compute response proportions (conditional probabilities) for each row of the contingency table (SI Fig. S1B). Psychophysical parameters (d’ and criteria) are related mathematically to each response proportion; the relationship is derived in previous work (Sridharan et al., 2014). Briefly, and as with the one-dimensional SDT model, higher sensitivity at a location typically yields a higher proportion of hits, a lower proportion of error responses (false alarms and misidentifications), or both. Higher criteria at a location typically yields a higher proportion of hits and false alarms.

To provide an intuition for how the sensitivities and criteria are related to the response proportions in the model, consider the Posner cueing task with two-alternatives (2-ADC task). In this model, the participant’s decision is modelled in a two-dimensional decision space with a bivariate decision variable (Ψ), modeled as a unit variance Gaussian random variable. The decision variable components for the cued and uncued locations – Ψ*_cued_* and Ψ*_uncued_*, respectively – are represented along the x- and y-axes respectively. On each trial, the values of these variables indicate the strength of the evidence for change at each location. The decision variable distribution for a “no change” event has a mean of zero and is centered at the origin (Fig. 1D, gray). For a change event at the cued or the uncued locations, the mean of the decision variable increases along the x-axis, or along the y-axis, respectively (Fig. 1D, cyan and magenta distributions). The mean of the signal distribution along each dimension, measured in units of the noise standard deviation for the respective dimension, quantifies perceptual sensitivity (d’) for detecting changes at the respective location.

To decide whether, and where, the change occurred, the observer adopts two decision threshold values, one for each location: *t_cued_* and *t_uncued_*. Based on these thresholds, the participant’s decision is modelled as follows: the participant reports a change at the cued location if Ψ*_cued_* > *t_cued_* and Ψ*_uncued_* < *t_uncued_*; the participant reports a change at the uncued location if Ψ*_uncued_* > *t_uncued_* and Ψ*_cued_* < *t_cued_*; the participiant reports “no change” if neither decision variable component exceeds its respective threshold. Lastly, if the decision variable components exceed their respective thresholds at both locations, the participant chooses the response for which the decision variable exceeds the threshold by a larger magnitude. Geometrically, this decision rule divides the space into three regions which correspond to the three alternatives that participants can choose from: change at the cued location, change at the uncued location or no change (Fig. 1D, cyan, magenta and gray lines).

The four m-ADC model parameters -- perceptual sensitivities at each location (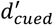 and 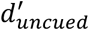) and thresholds at the two locations (*t_cued_* and *t_uncued_*) – are estimated from the response proportions in the 3x3 contingency table with maximum likelihood estimation (MLE) (Sridharan et al., 2014). From these values, SDT criteria (c) are computed as the magnitude of the threshold deviation from the equal likelihood (intersection) point of the signal and noise distributions, along the respective axis; in other words, *c* = *t* − *d′*/2 (Green & Swets, 1974). These criteria are inversely related to bias for reporting change at a location so that a lower value of the criterion (more liberal criterion) at a location reflects a higher bias for reporting a change at that location.

### EEG data acquisition and analysis

#### Data acquisition

Scalp EEG data were acquired with a EGI 128-electrode HydroCel EEG net (Magstim EGI, Eugene, Oregon, USA) connected to a Net Amps 400 amplifier (Magstim EGI, Eugene, Oregon, USA). The impedances of all the electrodes were maintained below 30 k**Ω** throughout the experiment (Barzegaran et al., 2012; Clementz et al., 2008; Friedl & Keil, 2021). Data was referenced to Cz during acquisition, digitized at 1000 Hz and stored for offline analyses.

We collected EEG data on all three experimental days, including the training session. To obtain robust Steady-State Visually Evoked Potentials (SSVEP) for each participant, we performed a titration experiment during the training session to select a pair of flicker frequencies that evoked prominent SSVEP spectral peaks in the two hemifields; these flicker frequencies ranged from 14-36 Hz (median 17 Hz) and were separated typically by 3-6 Hz across hemifields. For each participant, the pair of flicker frequencies were counterbalanced spatially, on the left and right hemifields, across blocks. Data were analyzed with a combination of Fieldtrip (Oostenveld et al., 2011), NoiseTools (de Cheveigné & Arzounian, 2018), and the Chronux toolbox (Bokil et al., 2010), as well as custom Matlab (2021a) scripts.

#### Preprocessing

EEG data were preprocessed with the FieldTrip toolbox (Oostenveld et al., 2011). Data from each task block for each participant were preprocessed separately. Data were downsampled to 250 Hz by decimation, re-referenced to the average of all the electrodes and demeaned. The data were then filtered in the range of [1, 100] Hz using a 4^th^ order Butterworth filter and subsequently with bandstop filters of the same order for the ranges [49, 51] Hz and [99, 101] Hz to remove line noise. The data were trial-epoched from fixation onset to response onset. We identified bad electrodes from the epoched data using the *nt_find_bad_channels* function (NoiseTools toolbox); a sample was flagged i) if its amplitude was greater than 3x the median absolute value of all the data or ii) if its amplitude was greater than 100 *μ*V. Data from an entire electrode was rejected if more than 50% of its samples were flagged. On average this resulted in <5% rejection rates; rejection rates in occipito-parietal cortex were typically <1%. Bad electrode traces were replaced with data from good electrodes using spherical spline interpolation (Oostenveld et al., 2011).

#### Rhythmic Entrainment Source Separation (RESS) for SSVEP quantification

In our experiment, two stimuli flickering at different frequencies were presented concurrently, one in each hemifield (see earlier Methods section on task description). To isolate SSVEPs corresponding to each flicker frequency with high signal-to-noise (SNR) we employed Rhythmic Entrainment Source Separation (RESS) (Cohen & Gulbinaite, 2017). For this, cross-channel covariance matrices (*C_S_,C_R_*) were computed based on data Gaussian filtered at the frequency of interest (signal, S) and other, reference frequencies (R) that are proximal to S. Here, we computed the signal covariance matrix (*C_S_*) by averaging covariance matrices of the data filtered in the first (mean: f_1_ Hz, width: 0.5 Hz) and second harmonics (mean: f_2_ Hz, width: 1 Hz) of each flicker frequency. The reference covariance matrix (*C_R_*) was constructed from the average covariance matrices of data filtered in the neighbouring frequencies of these harmonics (mean: f_1_±1 Hz, width: 1 Hz; mean: f_2_±2 Hz, width = 2 Hz). We applied generalized eigendecomposition of *C_S_* and *C_R_* 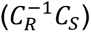 and chose the eigenvector corresponding to the highest eigenvalue, representing the subspace that maximized SSVEP power in the signal dimension relative to the reference dimension.

Because RESS provides more reliable components with longer data epochs, we constructed EEG epochs of 2 s duration, as follows: In our data, flickering stimulus presentation epochs were of variable lengths ranging from 650-3350 ms (350 ms prior to relevance cue onset followed by a variable post-relevance cue epoch ranging from 300-3000 ms). We grouped the trials into smaller groups pseudorandomly except for the condition that each group must have at least one trial longer than or equal to 2000 ms. Then EEG time traces in each group were averaged, after nan-padding shorter trials up to 2000ms. Because SSVEP phase is locked to the stimulus flicker phase signals, this operation had the added advantage of improving the signal-to-noise ratio of the evoked signal. Each of these averaged trials was divided into two overlapping sub-epochs comprising the initial 2000 ms and the final 2000 ms of each epoch. Following this, covariance matrix computation and generalized eigendecomposition were performed, as described above. Moreover, we constrained our SSVEP analysis to 58 parieto-occipital electrodes, which are known to typically evoke the strongest SSVEP oscillations (Chinchani et al., 2022). Denoised and preprocessed EEG traces were projected into the estimated RESS dimensions, and this data was used for all subsequent SSVEP analyses. RESS components were computed for each participant and task block separately, but after pooling trials across conditions within each block. The projected traces were then averaged across task blocks and frequencies for each participant and each condition.

#### Computing SSVEP power dynamics

To investigate stimulus-locked and change-locked trends in SSVEP power across conditions, we visualized the SSVEP power dynamics for each condition. For this analysis, we analyzed power at the second harmonic of the flicker frequency (2f), which has been reported to produce better lateralization and stronger attentional modulation (Kim et al, 2007); analysis with power at the fundamental (f), also produced qualitatively similar, albeit noisier results. To compute SSVEP power, first, we averaged RESS time traces for a given flicker frequency across trials to isolate the stimulus evoked periodic component for that frequency. We then employed multi-taper spectral estimation (time x half-bandwidth=1, number of tapers=1) (Bokil et al., 2010; McCoy et al., 1998) of the averaged RESS time course, using a sliding window of 500 ms duration sliding in steps of 20 ms. We computed SSVEP power dynamics: i) from 1000 ms before until 2000 ms after stimulus onset (stimulus-locked), and ii) from 1000 ms before until 1000 ms after the change epoch onset (change-locked); for plots in Figs. 3A-C, for each condition the power across both flicker frequencies were averaged. We tested if the SSVEP power modulations in these time windows, defined as [1000, 2000] ms after stimulus onset (stimulus-locked) and [-1050, -50] ms before change epoch onset (change-locked), differed significantly from zero using non-parametric Wilcoxon signed rank tests (Figs. 3A-C, insets).

To improve SNR further, we employed a jackknife procedure: in each window, SSVEP power was first computed for a jackknife replicate, by summing trial time courses leaving one trial (current trial) out, and then subtracted from the SSVEP power computed by summing all trial time courses. In practice, we observed that this approach mitigated uncorrelated noise across trials and limited noisy estimates of single trial SSVEP power. Tercile split analyses (see next) were performed on these jackknifed estimates of SSVEP power (Figs. 4A-D).

#### Analysis of induced spectrograms

Spectrograms were computed with a window of 500 ms duration sliding in steps of 40 ms, with multitaper spectral estimation (same parameters as mentioned before). Spectral power was divisively normalized on a per-frequency and per-electrode basis with a condition-specific baseline computed from -1000 ms to 0 ms before stimulus onset. As with the psychophysical parameters we computed power modulation with each type of cueing by taking the difference across cue validity levels (e.g., high relevance – low relevance), and performed cluster-based permutation tests to identify spectro-temporal clusters that showed significant cue-induced modulations (Figs. 3D-F, left panels, SI Figs. S4D-F). Although we report the cluster permutation test results, we computed alpha power time courses agnostic to these results; this was performed by averaging spectral power in a pre-defined, standard alpha frequency range (8-12 Hz) in a pre-defined window from 650-1650 ms after cue onset (or 1000-2000 ms after stimulus onset), with multitaper spectral estimation (same procedure as described above) (Figs. 3D-F right plots, SI Figs. S4D-F). We then computed a modulation index of alpha power, again, with a procedure similar to that described for SSVEP power. We tested if these alpha power modulations differed significantly from zero using non-parametric Wilcoxon signed rank tests (Figs. 3D-F, right panel, insets).

#### Tercile split analyses

Tercile split analyses based on neural signatures were performed for investigating the association between SSVEP (or alpha-band) power levels and modulations of psychophysical parameters. For example, for the tercile split analysis for d’ (or criterion), for each cue condition, we divided SSVEP power – computed from -1050 ms to -50 ms before change epoch onset – into three terciles, each containing 1/3^rd^ of the trials, in decreasing order of SSVEP power. We labeled the top third as “high SSVEP power” and bottom third as “low SSVEP power” trials. We computed contingency tables and estimated d’ (or criterion), separately, for these two sets of trials for each participant. We then compared whether d’ (or criterion) for high SSVEP power trials were different from d’ (or criterion) on low SSVEP power trials with a permutation test. We created a null distribution of sensitivity (or criterion) differences by swapping the low and high SSVEP labels for each participant independently 1000 times and compared the true difference to this distribution (Figs. 4A-D). An identical procedure was used for testing both d’ and criterion modulation based on a tercile split of pre-change (-1050 ms to -50 ms before change epoch onset) alpha-band power.

#### Neural markers of spatial expectation

We analyzed three recently reported EEG markers of expectation:

1. Pre-change alpha power: A previous study (Tarasi et al., 2022) reported that alpha power (8-12 Hz) was significantly suppressed contralateral to the high probability location compared to the low probability location. Because our tasks involved change detection we computed modulation index spectrograms locked to change onset similar to the ones computed locked to stimulus onset (see Methods, section on *Analysis of induced spectrograms*). Significance was assessed using cluster-based permutation tests (SI Figs. S4D-F).
2. Alpha phase opposition index: Another previous study reported that the phase opposition index – a measure of phase consistency for each response type (e.g., yes/no) – was significant in the alpha band (8-12 Hz) before the target event occurrence (Sherman et al., 2016). We followed a procedure closely similar to this study. The data were filtered in a 2 Hz band around each frequency of interest (f ± 2Hz). We then applied a Hilbert transform to obtain instantaneous phase and amplitude. For each frequency, the filter order was chosen such that the window length encompassed at least 4 cycles of the corresponding period. We then computed a measure of phase locking across trials by first normalizing the amplitude of the instantaneous analytic signal of each trial to 1.0 and then computing a complex mean vector across trials for every combination of time, frequency, condition (e.g., high or low probability) and response type. The magnitude of this complex mean vector was then averaged across responses and conditions to obtain the phase opposition values for each time and frequency (SI Figs. S4G-I), and applied a cluster-based permutation test to identify significant spectro-temporal clusters (see Methods, section on *Statistical Analysis*).
3. Fronto-parietal theta and alpha connectivity: Tarasi et al., 2022 also reported a significant increase in theta (5-8 Hz) connectivity contralateral to the high probability as compared to the low probability location (Tarasi et al., 2022). The connectivity was computed based on the weighted phase lag index (wPLI) (Vinck et al., 2011) between every pair of 8 predefined electrodes in frontal and parietal regions in two windows: post-relevance-cue ([0-400] ms after relevance cue onset, SI Fig. S4A), and pre-change ([-500, 0] ms from change epoch onset, SI Fig. S4B). The connectivity index (CI) was computed as the proportion of significant pairwise connections for each condition. Significance for the CI index was assessed with a random permutation test (see Methods, section on *Statistical Analyses*). Moreover, following more recent work (Di Gregorio et al., 2023), we also analyzed fronto-parietal alpha connectivity using a method similar to the theta connectivity, except that the data in the alpha (8-12 Hz) band was analyzed to compute the wPLI.

#### Electrodes used for analyses

For SSVEP analyses, we used all the parieto-occipital electrodes; these are shown in Fig. 3A. We used Rhythmic Entrainment Source Separation (RESS, (Cohen & Gulbinaite, 2017)) to obtain dimensions in this electrode space (a linear combination of electrodes) that maximally captured each stimulus’ flickering frequency. We then projected the data onto these dimensions and compared the SSVEP power between cueing conditions. For alpha power analyses, we considered posterior parieto-occipital electrodes symmetrically in both hemispheres; these are shown in Fig. 3D. For the fronto-parietal theta connectivity analysis, we provide the topoplots in SI Fig. S4C. These electrodes were chosen based on the electrodes on the EGI HydroCel-128 EEG electrode cap closest to the ones used in (Tarasi et al., 2022).

### Neural representations of relevance, probability and Posner cueing

#### i) Representational Similarity Analysis (RSA)

We employed Representational Similarity Analysis(Sheahan et al., 2021) to characterize the nature of neural (EEG) representations associated with each type of cueing (relevance, probability) in the dual cueing task. We constructed a representational similarity (or dissimilarity) matrix (RDM) for the EEG data (RDM_EEG_) by computing a dissimilarity metric between every pair of conditions. EEG data were first z-scored over all trials for each electrode and time point and averaged condition-wise. Then, we performed a regression across trials for each condition and electrode. Each regressor was a binary vector with a value of 1 if the trial included the specific combination of conditions (one of 66) or 0 otherwise. Regression weights for each electrode were normalized by the residual covariance matrix; this procedure has been previously employed to improve signal-to-noise of the weight estimates (Walther et al., 2016). These normalized weight vectors were then correlated across every pair of conditions, and the correlation coefficient was subtracted from 1, to yield the neural RDM.

We specified 3 different binarized model RDMs: i) a relevance RDM (high/+1 or low/-1 relevance) (Fig. 5A, model RDM for *β_relevance_*), ii) a probability RDM (high/+1 or low/-1 probability) (Fig. 5A, model RDM for *β_probability_*) and iii) a visual RDM based on the distances between mean orientations of the gratings, divided into 3 orientation bins (Fig. 5A, model RDM for *β_visual_*). This yielded 66 (^12^C_2_) combinations of conditions per RDM matrix. After constructing these matrices, we z-scored them. EEG RDMs were computed at each timepoint separately and regressed against model RDMs as follows.

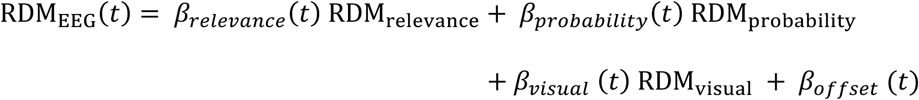

Following this regression, the timecourse of each regression weight was plotted as a function of time (e.g., Fig. 5B). Higher values of a weight (e.g., *β_relevance_*) at a particular timepoint indicate greater information in the EEG signal about the corresponding condition (e.g., attention) at that timepoint, reflecting better separation of EEG neural representations across different levels of that condition (e.g., high versus low relevance). β coefficients were computed separately for each block and each side (left, right) with 59 contralateral electrodes in each hemisphere (all except the midline electrodes; Fig. 5A); results were then averaged across blocks and hemispheres.

#### ii) Decoding cue locations with embedding-aided deep convolutional neural networks (CNNs)

To characterize neural representations associated with each type of cueing we developed a novel “embedding-aided” deep convolutional neural network architecture (Biswas et al., 2023) to predict cue location based on single trial EEG spectrogram tensors (see next). Independent models were trained for predicting the relevance cue location (“Attention-CNN”), probability cue location (“Expectation-CNN”) or the Posner cue location (“Posner-CNN”).

#### Data Preprocessing

Data were epoched from 1350 ms before to 1000 ms after relevance cue onset; this included a pre-stimulus epoch of 1000 ms that followed probability cue onset. Trials with cue durations <1000 ms, were padded zeros; a Kolmogorov-Smirnov test revealed that cue duration distributions were not statistically significant different across cue locations (p>0.97 for all cue types). Spectrograms were computed in sliding windows of 500 ms window (shift: 20 ms) with multitaper spectral estimation as described in the sections on *Analysis of induced spectrograms, computing SSVEP power dynamics*. Frequencies were sampled from 3-45 Hz with a 2 Hz resolution. This yielded single-trial spectrogram tensors of 93x21x128 (time x frequency x electrodes) dimension, which were provided as input to the CNN.

#### Model Architecture

Initially, the input tensor was convolved with learnable filters from two “embedding” layers. First, we designed an “electrode embedding” layer (Fig. 6A. “elec. emb.”) to account for the variability in positioning of electrodes across participants and experimental blocks relative to putatively homogeneous underlying neural sources. This layer comprised filters of dimension (1x1x128)x128, i.e., 128 – 1x1x128 filters. Convolving these filters with the input spectrogram, resulted in the signal from each channel being replaced by a linear combination of signals from all other channels. Second, we designed a “time-frequency” embedding layer (Fig. 6A, “t x f emb.”) to address variations in spectral peaks and signal latency across participants and experimental blocks. In this case, the output was convolved depth-wise with a learnable filter of dimension (3x3x128) (Chollet, 2017).

The preprocessed spectrograms of dimension - 93x21x128, were provided as input to a deep CNN model whose weights were common across participants and experimental blocks. This model contained 3 serial convolution layers, each with filters of dimension (7x7x128)x96, (5x5x96)x64, and (3x3x64)x32 respectively. The strides along both time and frequency dimensions were set to 1, and the input was padded to conserve the dimension after convolution. Each convolutional layer was followed by a “max-pool” layer (dimensions: (3x2), (3x2) and (2x2), respectively). Strides matched the filter dimensions in these layers. The output of this CNN was flattened to obtain a 320-dimensional vector which was followed by a fully connected layer. The output of this layer (a 40-dimensional vector) was concatenated with a 5-dimensional participant-specific embedding vector. This embedding vector was obtained from a “participant”-embedding layer which accounts for miscellaneous sources of neural variability arising from variations in behavioral state (e.g., alertness, motivation) across participants. The resulting 45-dimensional feature vector was provided as input to a binary classifier to predict cue locations (left or right).

The model was trained using Adam optimization with a batch size of 32 and an inverse-time learning rate decay (starting value: 10^-4^). Based on empirical observations of the learning curve (rate of decrease in binary cross entropy error), the number of training epochs was fixed at 15. Model hyperparameters, including the L2 regularization coefficient, and dropout were tuned using the Tree-structured Parzen Estimator (TPE) algorithm (Bergstra et al., 2011) to maximize accuracy on a held-out set (20% of training data).

#### Saliency Maps

We computed saliency maps (Simonyan et al., 2013) to identify neural (spectro-temporal) features most predictive of each type of cue location. For each CNN, saliency maps *S*(*X*) were computed as:

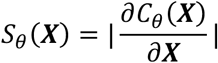

where ***X*** denotes the input vector and *C_θ_*(***X***) denotes the predicted class. The absolute value reflects the fact that the magnitude of the gradient, and not its sign, is relevant as measure of the saliency. Saliency maps were computed for all three CNNs (Attention, Expectation and Posner). independently.

#### Comparing saliency maps

To compare saliency maps of Attention-CNN and Expectation-CNNs with those of the Posner-CNN, we computed two measures – Mean Absolute Error (MAE) and pairwise “robust” correlations between their respective saliency maps. We compared the saliency map values in the post-relevance cue window ([600-1100] ms post-relevance cue) for the Attention-CNN and post-probability cue window ([250-750] ms post-probability cue) for the Expecatation-CNN (Fig. 7A,B) with the post-cue window for Posner-CNN ([600-1100]ms post-cue). For computing the Mean Absolute Error (MAE), for each participant, we vectorized the saliency values in the aforementioned windows and computed the MAE according to the following equation:

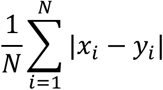

where *x_i_* and *y_i_* are vectorized saliency map values of the corresponding CNN pairs, and N is the number of values in each of the vectorized saliency maps. For computing the robust correlation, we computed the bend correlation between the pariwise vectorized saliency values in the same, aforementioned windows.

### Statistical analyses

To test whether each type of cue modulated psychophysical parameters (d’ and c) significantly we employed a Wilcoxon signed rank test. We compared the psychophysical parameters between high versus low relevance conditions, high vesus low probability conditions and Posner cued versus uncued conditions. To test for interaction effects between attention and expectation on psychophysical parameters, we performed an ANOVA on d’ and c separately with relevance cue level (high, low), probability cue level (high, low) as factors and participants as random effects followed by a post-hoc Tukey HSD tests to identify significant pairwise differences. For correlations between parameter modulations across tasks, we report the significance value obtained with a percentage bend correlation (bend criterion: 0.2) (Wilcox, 1994). To compare magnitudes of sensitivity or criterion correlations across tasks, we used the Pearson and Filon’s z test from the “compcorr” toolbox (Diedenhofen & Musch, 2015).

To compare modulations in SSVEP power timecourses across cue levels (high, low) for the three types of cueing, we performed a cluster-based permutation test (Maris & Oostenveld, 2007). To compare the SSVEP power (|S|) in post-relevance cue and pre-change windows, we computed modulation indices (MI=(|S|_high_-|S|_low_)/(|S|_high_+|S|_low_)) and performed a Wilcoxon signed-rank test on these modulation indices to test for significant difference from zero.

To identify behavioral correlates of SSVEP power modulation, we performed a tercile split analysis on the cued SSVEP power for the high relevance and high probability conditions. To statistically test if the psychophysical parameter for the two terciles were significantly different, we performed a one-tailed permutation test by shuffling the high and low SSVEP power labels for the respective behavioral parameter, randomly for each participant, and computed the difference in the parameter values between these conditions. We repeated this shuffling procedure 1000 times with different random seeds to create a null distribution of difference values and compared the true difference against the null distribution.

To test for significant cue-induced modulation in the time-frequency spectrograms, we performed cluster-based permutation tests on difference spectrograms locked to stimulus and change epoch onsets. To test for significant alpha-band (8-12 Hz) lateralization, we averaged the power in the alpha frequencies and performed cluster-based permutation tests between cue levels (high versus low) on the alpha power timecourses locked to stimulus and change. As with the SSVEP analysis, we performed Wilcoxon signed-rank tests on modulation indices computed for predefined post-relevance cue and pre-change windows.

Significance testing for wPLI analysis was performed by first computing a connectivity index (CI). First, a one-tailed Wilcoxon signed-rank test (p_threshold_=0.05) was used to identify significant connections with higher wPLI values for the high, compared to the, low probability condition; the CI represents the fraction of all significant connections. To test for significance, the a null distribution of CI values was computed by swapping the high and low probability labels 1000 times randomly across the participants. The p-value was computed as the proportion of null distribution CI values greater than the actual CI.

For the phase opposition index analysis, for each of the time, frequency and condition, we created a surrogate distribution by first randomizing the “yes” and “no” labels across trials keeping the number of trials for each label constant. A null distribution of phase opposition measures was computed by repeating this procedure 500 times for each time and frequency value. p-values were computed by comparing the actual phase opposition index against the null distribution value. These p-value maps were averaged across conditions, and cluster-based permutation tests were then used to identify significant spectro-temporal clusters (SI Figs. S4G-I).

In the decoding analysis with deep CNNs, median and 99% confidence intervals across participant are report for each self-decoding analysis; for each participant and decoding condition accuracies were averaged across 25 test sets (5 left-out folds x 5 seeds). Decoding performance in which the lower limit of the 99% CI fell above 50% were considered to be significantly above chance (Fig. 6B). Correlations between average saliency map values between the Posner-CNN and the Attention- or Expectation-CNNs were assessed in predefined time and frequency windows with robust correlations (Figs. 7D-F)

## Data Availability

The data and code necessary to reproduce the key results of the study is available at the OSF repository: https://osf.io/gjwy2/?view_only=9af0f40e549249ee9cde431c86137ddf

## Acknowledgments

This research was supported by a Department of Biotechnology-Wellcome Trust India Alliance Intermediate fellowship (grant number IA/I/15/2/502089), a Department of Science and Technology Swarna Jayanti Fellowship (grant number SB/SJF/2021-22/02), a Pratiksha Trust Intramural grant (grant number KVCH22-2047), and an India-Trento Program for Advanced Research grant (grant number INT/ITALY/ITPAR-IV/COG/2018/G)

