## Supplementary Information for "Neural mechanisms of attention, not expectation, govern spatial selection by probabilistic cueing"

### Supplementary Results

#### *Neural markers of spatial expectation*

Recent work suggests that increase in pre-stimulus (anticipatory) fronto-parietal theta connectivity and a decrease in pre-stimulus alpha power (Tarasi et al., 2022; van Ede et al., 2020) in the occipital electrodes may index higher signal expectation, but the latter marker has not been consistently replicated (Zhou et al., 2021). In addition, a phase opposition index in the alpha band, which measures the phase locking of band-limited event traces across trials(Sherman et al., 2016), has been found to differentially predict “Yes” and “No” responses. We tested for the modulation of these three neural markers – fronto-parietal theta connectivity, alpha power modulation and alpha phase opposition index – by all three types of cueing (see Methods, section on *Neural markers of spatial expectation*). We examined these signatures in a pre-change (-1050 to -50 ms before change) epoch rather than the pre-stimulus epoch because orientation change was the relevant event of interest for our tasks.

Interestingly, fronto-parietal connectivity index ($C_{ind}$) in the pre-change epoch was weakly modulated by Posner cueing ($p=0.018$, random permutation test) (SI Figs. S4A-B), but not by the other types of cueing ($p=0.450$, and $p=0.617$ for relevance and probability cueing, respectively). Moreover, we observed that alpha power was systematically suppressed in the pre-change epoch both by relevance cueing and Posner cueing (SI Figs. S4D, F) ($p<0.05$, cluster permutation test), but not by probability cueing (SI Fig. S4E), mimicking the effects observed in the post-cue epoch. Finally, we did not observe any significant change in the phase opposition (PO) index in the alpha range prior to change onset, with any type of cueing (no clusters with $p < 0.05$ for any type of cueing; cluster permutation test) (SI Figs. S4G-I). Furthermore, following more recent work, we also analyzed fronto-parietal alpha connectivity as an index of post-stimulus processing (Di Gregorio et al., 2023) (Methods), but found no significant modulation of this connectivity either in the post-relevance-cue epoch ($p>0.15$) or the pre-change epoch ($p>0.2$).

### Supplementary Figures and Tables

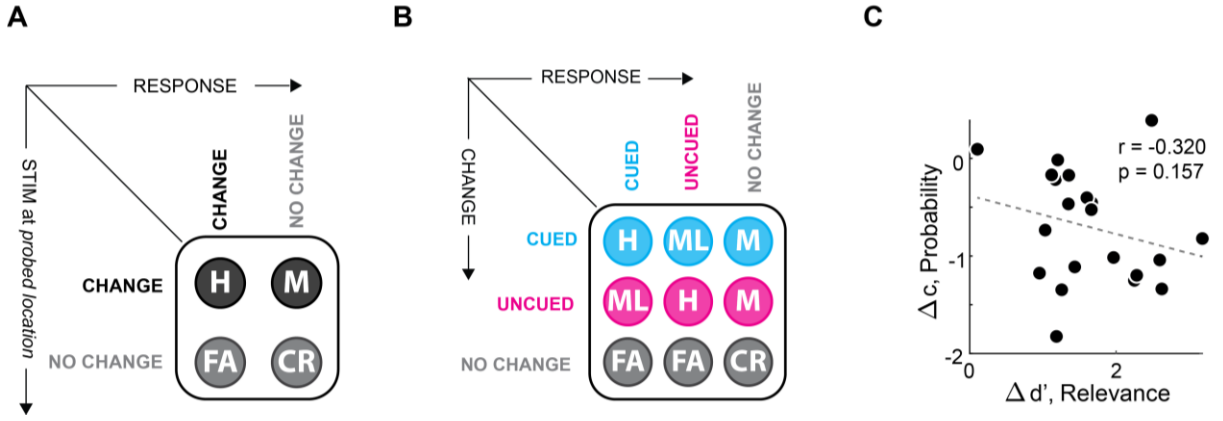


**SI Figure S1. Contingency tables and additional results on psychophysical analysis.**

- **A.** Stimulus-response contingency table in the dual cueing task. (Rows) Potential stimulus events at the probed location (change/no change). (Columns) Potential response types (Change/No Change). H: hits, M: Misses, FA: False Alarms, CR: Correct Rejections. Four such tables were constructed, one for each combination of relevance and probability cueing levels (see Figure 1A main text, inset).
- **B.** Stimulus-response contingency table in the Posner cueing task. (Rows) Potential stimulus events (change at cued location, uncued location or no change). (Columns) Potential response types (cued change, uncued or no change.
- **C.** Same as in Figure 2D (main text) but showing the correlation between sensitivity (d') modulation by relevance cueing (x-axis) and criterion (c) modulation by probability cueing (y-axis) in the dual cueing task. Other conventions are the same as in Figure 2D (main text).

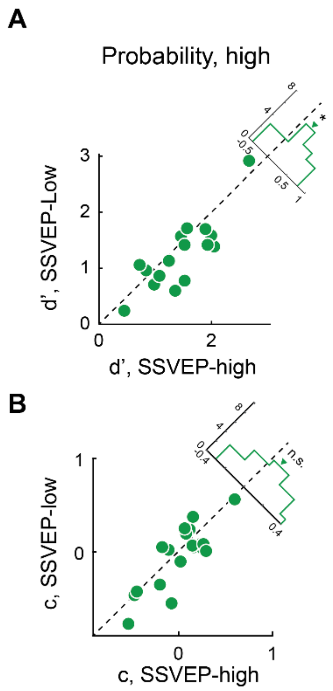


**SI Figure S2. Additional results on SSVEP power analysis.**

- **A.** Same as in Figure 4A (main text) but showing sensitivities (d') based on a tercile split of SSVEP power for the high probability condition in the dual cueing task. Other conventions same as in Figure 4A (main text).
- **B.** Same as in Figure 4B (main text) but showing criteria (c) based on a tercile split of SSVEP power for the high probability condition in the dual cueing task. Other conventions same as in Figure 4B (main text).

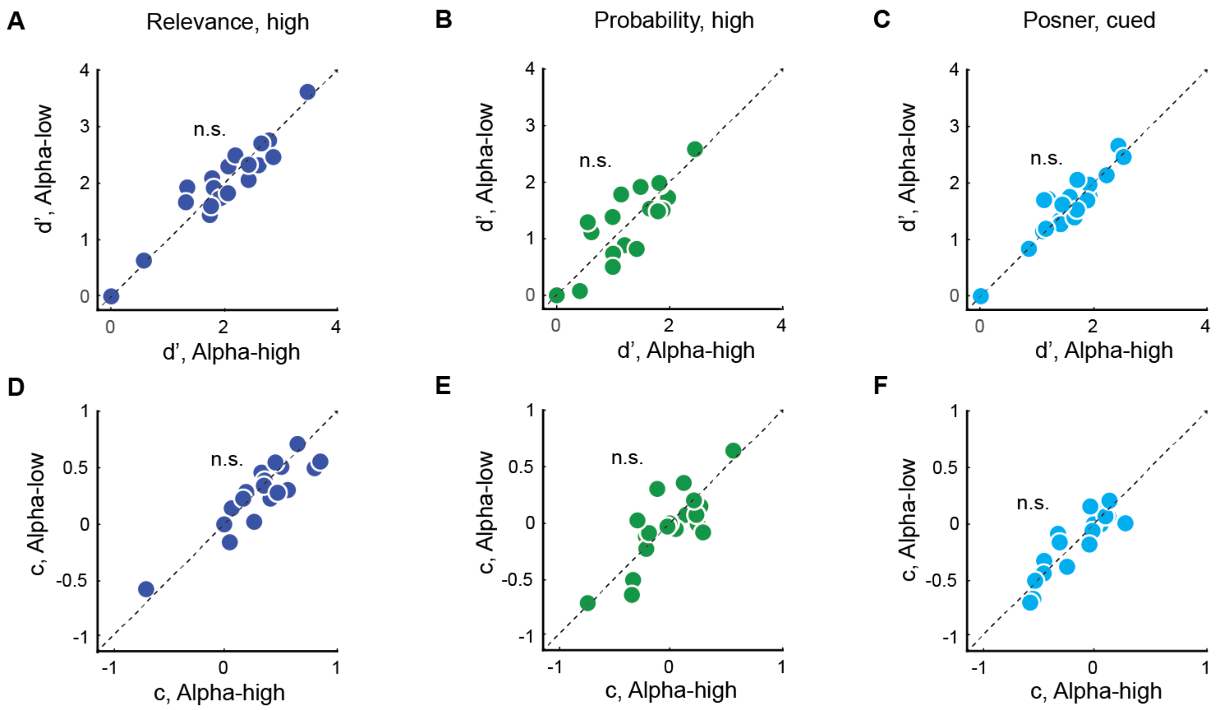


**SI Figure S3. Pre-change alpha tercile split analysis**

**A-C.** Same as in Figure 4A (main text) but showing sensitivities (d') based on a tercile split of change-locked alpha power in the dual cueing task, for the high relevance (panel A), high probability (panel B) and Posner cued conditions (panel C), respectively.

**D-F.** Same as in Figure 4B (main text) but showing criteria (c) based on a tercile split of change-locked alpha power in the dual cueing task, for the high relevance (panel D), high probability (panel E) and Posner cued conditions (panel F), respectively.

(A-C) Other conventions are the same as in main Figure 4A.

(D-F) Other conventions are the same as in main Figure 4B.

low cue level; gray bar: low > high cue level. Error bars: jackknife standard error of mean. *p<0.05.

**B.** Same as A but showing wPLI computed in a 500 ms period before change epoch onset. Other conventions are the same as in panel A.

**C.** Frontal (red) and parietal (blue) electrodes from which the connectivity index was computed, for trials in which the right hemifield was analyzed (yellow rectangle). The corresponding mirror-symmetric configuration of electrodes was chosen for trials where the left hemifield was analyzed.

**D.** Same as Figure 3D (left, main text) but showing broadband spectral modulation by relevance cueing, locked to change epoch onset (black vertical line).

**E.** Same as in panel D, but showing broadband spectral modulation by probability cueing, locked to change epoch onset. Other conventions are as in panel D.

**F.** Same as in panel D, but showing broadband spectral modulation by Posner cueing, locked to change epoch onset. Other conventions are as in panel D.

**G.** Spectrotemporal map of the phase opposition index values for relevance cueing (average across cue levels) in the dual cueing task (see Methods, section on Neural markers of spatial expectation: Alpha phase opposition index). x-axis: time from change onset (white vertical line); y-axis: frequency; z-axis: p-values. Cooler colors reflect higher significance levels.

**H.** Same as in panel G but showing phase opposition index values for probability cueing (average across cue levels). Other conventions are the same as in panel G.

**I.** Same as in panel G but showing phase opposition index values for Posner cueing (average across cue levels). Other conventions are the same as in panel G.

**SI Table S1. CNN model hyperparameters.** Hyperparameters used while training the deep convolutional neural networks. These hyperparameters were identical across the Attention-CNN, Expectation-CNN and Posner-CNN.

| **Hyperparameter** | **Value** |
|---|---|
| L2 regularization coefficient for embeddings | 0.0031 |
| L2 regularization coefficient for dense layers | 0.022 |
| L2 regularization coefficient for convolutions | 0.0035 |
| Dropout for the dense layer weights | 0.34 |
